# Genome-wide analysis reveals the importance of histone acetyltransferase Esa1 in transcriptional regulation during nitrogen starvation

**DOI:** 10.64898/2026.07.29.741553

**Authors:** Uzair Khan, Krystal Cvetkovski, Matthew Werick, Cole Tracy, Sara Fatima, Dimitra Dialynaki, Daniel J. Klionsky, Chhabi K. Govind

## Abstract

Macroautophagy/autophagy is a process that degrades intracellular components and is strongly triggered by nitrogen starvation (-N). Some *ATG* (autophagy related) genes are activated at the transcriptional level in nitrogen starvation; however, a full understanding of transcriptional induction and the role of chromatin during this process remains unclear. To address this, we measured the occupancy of RNA polymerase II (Pol II), histone H3, and acetylated H4 (H4Ac) under nutrient-rich and -N conditions by ChIP-seq. We found that most genes are rapidly downregulated within 15-30 min, including ribosomal protein (RP) and biogenesis (RiBi) genes. Meanwhile, genes involved in amino acid (AA) biosynthesis are upregulated, along with many *ATG* genes. Unexpectedly, RP and RiBi genes were reinduced by 3 hours. Furthermore, many upregulated genes remained active during prolonged starvation. Histones are typically removed from promoters during transcription activation. Consistent with this, we found that most induced genes exhibited histone eviction and increased H4 acetylation at their promoters, suggesting a possible role for histone acetylation in their activation. In line with this, depleting Esa1, an essential H4 histone acetyltransferase, nearly abolished the induction of ribosomal biosynthetic genes and many AA biosynthetic genes. Sustained activation of many genes during prolonged starvation highlights the vital role of transcription in supporting autophagy and cell survival. This is the first comprehensive study to detail changes in chromatin, histone acetylation, and transcription during nitrogen starvation, highlighting the importance of Esa1 and H4Ac in this process.

## Introduction

Autophagy is a conserved cellular process that degrades and recycles cellular components, including proteins and organelles [1]. Selective autophagy is utilized to degrade specific cargo, such as ribosomes via ribophagy or mitochondria through mitophagy [2–5]. Whereas autophagy occurs at a basal level, it is significantly induced by various stresses and is also utilized for the degradation of specific cargo [5–9]. It plays a particularly crucial role during starvation stress by degrading cellular components and replenishing precursors necessary for continued metabolism. Inhibiting the TOR pathway through rapamycin or nutrient deprivation triggers autophagy [3, 10–12]. In yeast, autophagy is potently induced by nitrogen starvation [13, 14] and it is regulated at the transcriptional, translational, and post-translational levels [15, 16].

Several *ATG* (autophagy related) genes have been identified, some of which participate in selective autophagy [17–20]. A few of the *ATG* genes are transcriptionally upregulated when cells are subjected to nitrogen starvation. Additionally, many other factors, including transcription activators, repressors, histone-modifying enzymes, and chromatin remodelers, contribute to autophagy in yeast. For example, the binding of Ume1 to the *ATG8* promoter represses its transcription [21], potentially by recruiting the Rpd3 histone deacetylase complex (HDAC), which deacetylates histones to repress transcription [22–24]. In line with this, the loss of Rpd3 enhances autophagy, suggesting that Rpd3 acts as a negative regulator of this pathway [25]. Rpd3 also deacetylates the chromatin remodeler ATPase Ino80 and H2A histone variant H2A.Z [26]. In contrast, Esa1, an essential histone acetyltransferase (HAT), promotes autophagy by acetylating Atg3 [27]. Thus, both the H4 HAT and HDACs regulate autophagy in a contrasting manner.

Esa1 is the catalytic subunit of the NuA4 HAT complex [28] and is the only enzyme that acetylates the H4 amino-terminal tail lysine residues in *S. cerevisiae* [29–31]. It is recruited to the promoters of ribosomal protein genes (RPGs), leading to increased histone H4 acetylation (H4Ac) [31]. However, recent studies have shown that although Esa1 acetylates nucleosomes at RP gene promoters, its loss only modestly reduces transcription of RP genes under nutrient-rich conditions [32]. Given that Esa1 is the sole H4 HAT in yeast and that it promotes autophagy [27], it remains to be seen whether Esa1 enhances autophagy by stimulating transcription of specific genes in -N conditions, in addition to acetylating Atg3.

Most research on the regulation of autophagy has focused on the expression of specific *ATG* genes [15, 33]. Our understanding of cellular responses to nitrogen starvation, a potent inducer of autophagy, particularly regarding the timing of gene expression during this phase, remains limited. During starvation, autophagy is vital for recycling cellular components and aids in replenishing necessary proteins, enzymes, and other essential factors to alleviate the effects of starvation. For instance, degradation of proteins by autophagy helps in increasing the amino acid (AA) pool in nitrogen-starved cells [34]. However, nitrogen starvation also increases expression of the transcriptional factor Gcn4, which could transcriptionally induce AA biosynthetic genes, thereby obviating the need for autophagy. A recent study revealed that several AAs are synthesized during autophagy, and recycled AAs are channeled to pathways leading to glutamate synthesis [35]. However, what kind of transcriptional response is mounted by cells during nitrogen starvation remains to be determined.

In this study, we determined global changes in transcription in response to short-term and long-term nitrogen starvation (-N). We employed RNA polymerase II (Pol II) chromatin immunoprecipitation sequencing (ChIP-seq) to evaluate precise changes in transcription. Additionally, we determined genome-wide levels of H3 and H4Ac under starvation conditions. These global analyses revealed that both ribosomal protein (RP) and biogenesis (RiBi) genes, along with many other genes, are immediately downregulated upon sensing nitrogen starvation. At the same time, cells significantly upregulate the transcription of genes involved in AA synthesis. Notably, these genes involved in ribosome biosynthesis are reactivated to levels comparable to those seen in nutrient-rich conditions within 3 h of N starvation. These changes in global transcription coincide with changes in histone and acetylated histone occupancies, suggesting that chromatin remodeling may be necessary for transcription during induction. Consistent with this, depleting the H4 HAT Esa1 nearly abolished the transcriptional reinduction of RP genes and severely reduced the transcription of AA biosynthesis genes, illustrating the importance of Esa1 in promoting gene expression under autophagy-inducing conditions.

## Results

### Nitrogen starvation triggers global transcriptional downregulation

To examine the dynamic changes in transcription in response to nitrogen starvation, we performed ChIP-seq for Rpb3 (a Pol II subunit) in cells grown in nutrient-rich medium (YPD) and in cells subjected to N starvation at specific time points. Chromatin was isolated from cells cultured in YPD and from cells starved for 0, 5, 15, and 30 min; 1, 3, 6, 18 h; and 1, 2, 3, 4, and 7 d. Pol II occupancy was calculated at each base pair and averaged across each open reading frame (ORF).

Because N starvation triggers autophagy, we expected cells to lower the expression of non-essential genes while increasing the expression of genes crucial for coping with starvation, including *ATG* genes. The ChIP-seq data show that Pol II occupancy was significantly downregulated as early as 5 min into starvation, with a substantial reduction observed at the 1-h time point (Fig. 1A and Table S2). Thus, nitrogen deprivation causes a rapid decrease in transcription, as shown by the immediate reductions in Pol II occupancies after transferring cells to -N medium (Fig. 1A). We noted that the trendline for the 3-h time point was higher than that seen at the 30-min time point (Fig. 1A, compare dashed blue [30-min time point] with dashed green [3-h time point] lines), suggesting potential transcriptional induction of some genes following initial downregulation.

**Figure 1.**
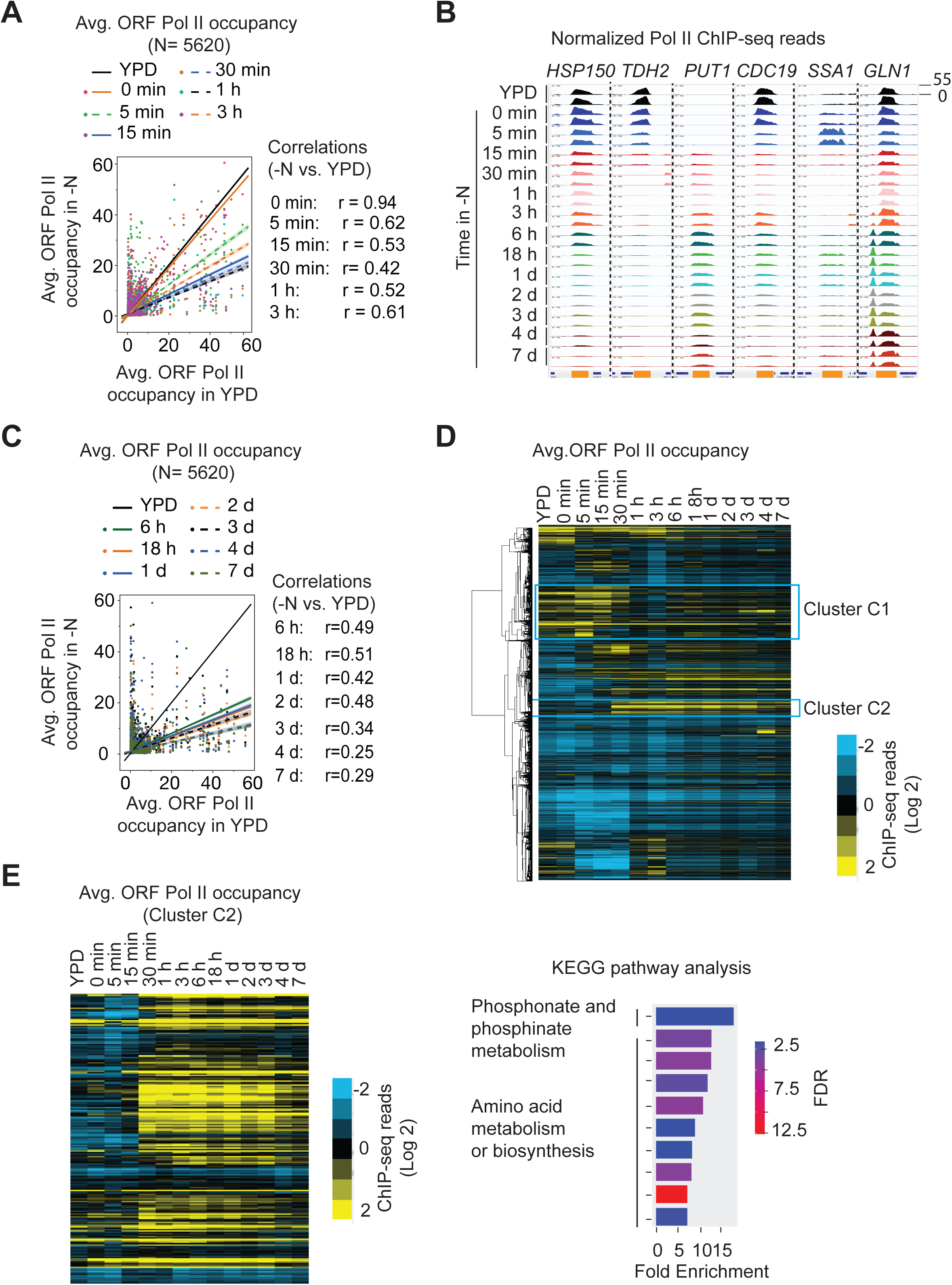
Most genes are rapidly downregulated in nitrogen-starvation conditions. Cells were either grown in YPD or transferred to N-starvation media (-N) for different time periods and subjected to RNA polymerase II (Pol II) ChIP-seq. Pol II ChIP-seq reads were normalized, and the average read count was calculated for each ORF, referred to as average (avg.) ORF Pol II occupancy. (**A**) Scatter plot showing Pol II occupancy for 5620 genes in YPD and in -N for specified time periods (0-min to 3-h time points). The black line indicates no change in Pol II occupancy (YPD). Pearson correlation is shown for each time point relative to YPD on the right-hand side of the plot. (**B**) Genome browser screenshot showing normalized Pol II ChIP-seq reads in YPD and -N from 0- min to 7-d time points. The reads for each time point are displayed in different colors, and the replicates are shown in the same color. The height indicates the number of reads in the plotted area (scale: 0–55). A vertical dashed line separates the non- contiguous genes shown here. (**C**) Scatter plot showing Pol II occupancies in YPD and at longer time points in -N conditions (6-h to 7-d). (**D**) Hierarchical clustering of Pol II occupancies in YPD and -N is depicted as heatmaps for all yeast genes. Two distinct clusters are labeled C1 and C2. (**E-F**) The heatmap for the C2 cluster (in panel D) is zoomed in, and the gene-set enrichment for the C2 genes is shown in the right-hand panel. False discovery rates (FDRs) are computed via the Benjamini-Hochberg method to correct for multiple testing. Fold enrichment is defined as the percentage of genes in a pathway divided by the corresponding percentage in the background.

We observed several transcriptional induction trends. For instance, *PUT1*, which encodes proline oxidase, was induced within 15 min of starvation, and it continued to be transcribed throughout the starvation period (Fig. 1B). This makes sense because Put1 is involved in utilizing proline as the sole nitrogen source. Previous studies have noted that the genes in glycolytic pathways are downregulated when TOR is inhibited [36]. Consistent with this, enzymes involved in glycolysis, *TDH2* and *CDC19,* were quickly downregulated. However, we observed a slight reinduction of CDC19 starting at 3 hours, followed by a gradual decline thereafter. *SSA1*, encoding an ATPase that assists in protein folding, exhibited a brief spike at 5 min, likely due to starvation stress, but was subsequently downregulated. Ssa1 activates the cAMP-PKA pathway, which suppresses growth and promotes cell entry into the stationary phase [37, 38]. While *HPS150* exhibited slower downregulation kinetics, Pol II occupancy remained high, aligning with studies indicating that this gene is regulated by both heat shock and nitrogen starvation [39]. To determine whether transcriptional upregulation continues, we analyzed Pol II occupancies at later times, including a 7-d time point. In general, Pol II occupancy decreased over time, reaching its lowest point for most genes on day 7 (Fig. 1C; the last time point analyzed). The data indicate that N starvation caused widespread transcriptional downregulation that persisted throughout the starvation period. However, some genes, such as *PUT1,* were induced, while others remained active throughout the starvation period. For example, *GLN1*, involved in glutamine synthesis, remained activated throughout the starvation period analyzed in the current study (Fig. 1B).

To gain insights into how nitrogen starvation affects the expression of genes typically transcribed under nutrient-rich conditions, we focused on the top 25% of genes exhibiting the highest average Pol II occupancy in their coding regions (T25; N=1405) in YPD conditions. Of the T25 genes, 414 were found to be downregulated by more than 2-fold after 30 min (Fig. S1A). These genes displayed significant reductions within just 5 min of starvation, indicating that the downregulation was almost instantaneous. Many of the downregulated genes are involved in nucleotide metabolism and glycolysis/gluconeogenesis (Fig. S1B). Additionally, genes related to ribosome biogenesis also showed reduced transcription by the 30-min time point (analyzed separately in Fig. 3). In contrast, 374 genes exhibited a less than 25% change in Pol II occupancy at the 30-min time point. The relatively unchanged transcription of these genes suggests that they may continue to play a key role in the stress response. Notably, these genes showed a slight upregulation at 15-30 min before returning to the YPD level of Pol II in their ORFs. Some of these unchanged genes are associated with the phagosome, proteasome, and endocytic processes (Fig. S1C-S1D).

Hierarchical clustering of genes, excluding RP genes, revealed that while most genes were downregulated, several were upregulated, with some exhibiting transient upregulation during starvation (Fig. 1B and 1D). The genes in cluster 1 (C1; Fig. 1D) highlight the genes expressed in YPD that were subsequently downregulated. Gene ontology (GO) analyses revealed that these genes are involved in oxidative phosphorylation and nucleotide biosynthesis. One of the clusters (C2) contained genes that remained consistently upregulated throughout most of the starvation period (Fig. 1D and 1E). This cluster included genes involved in amino acid biosynthesis and metabolism. In summary, the Pol II ChIP-seq data revealed that nitrogen starvation induces an immediate, genome-wide reduction in transcription.

### A significant number of ATG genes are transcriptionally induced during nitrogen starvation

Autophagy is induced during nitrogen starvation [40]; however, most studies have only examined autophagy induction at intermediate time points such as between 1- and 4-h of starvation. To monitor autophagy induction, we used the GFP-Atg8 processing assay [41]. Atg8 is conjugated to phosphatidylethanolamine on both sides of the membrane of the initial sequestering compartment of autophagy, the phagophore. Upon autophagosome maturation, the Atg8 on the outer surface is removed (deconjugated) and reused, whereas the population on the inner side of the autophagosome remains within this compartment and is delivered to the vacuole along with the cargo. Thus, Atg8 is the only yeast Atg protein that remains associated, at least to any substantial extent, with the completed autophagosome. The GFP-Atg8 chimera follows the same transport route; however, GFP is much more resistant to proteolytic degradation than Atg8, resulting in the generation of free GFP within the vacuole lumen. Accordingly, the amount of free GFP in cells expressing GFP-Atg8 provides an indication of autophagic flux. We observed the induction of autophagy under conditions of nitrogen starvation, as evidenced by the increased accumulation of free GFP in cells starved for nitrogen for 60 min compared to cells in YPD or -N media for 0 min (Fig. 2A).

**Figure 2.**
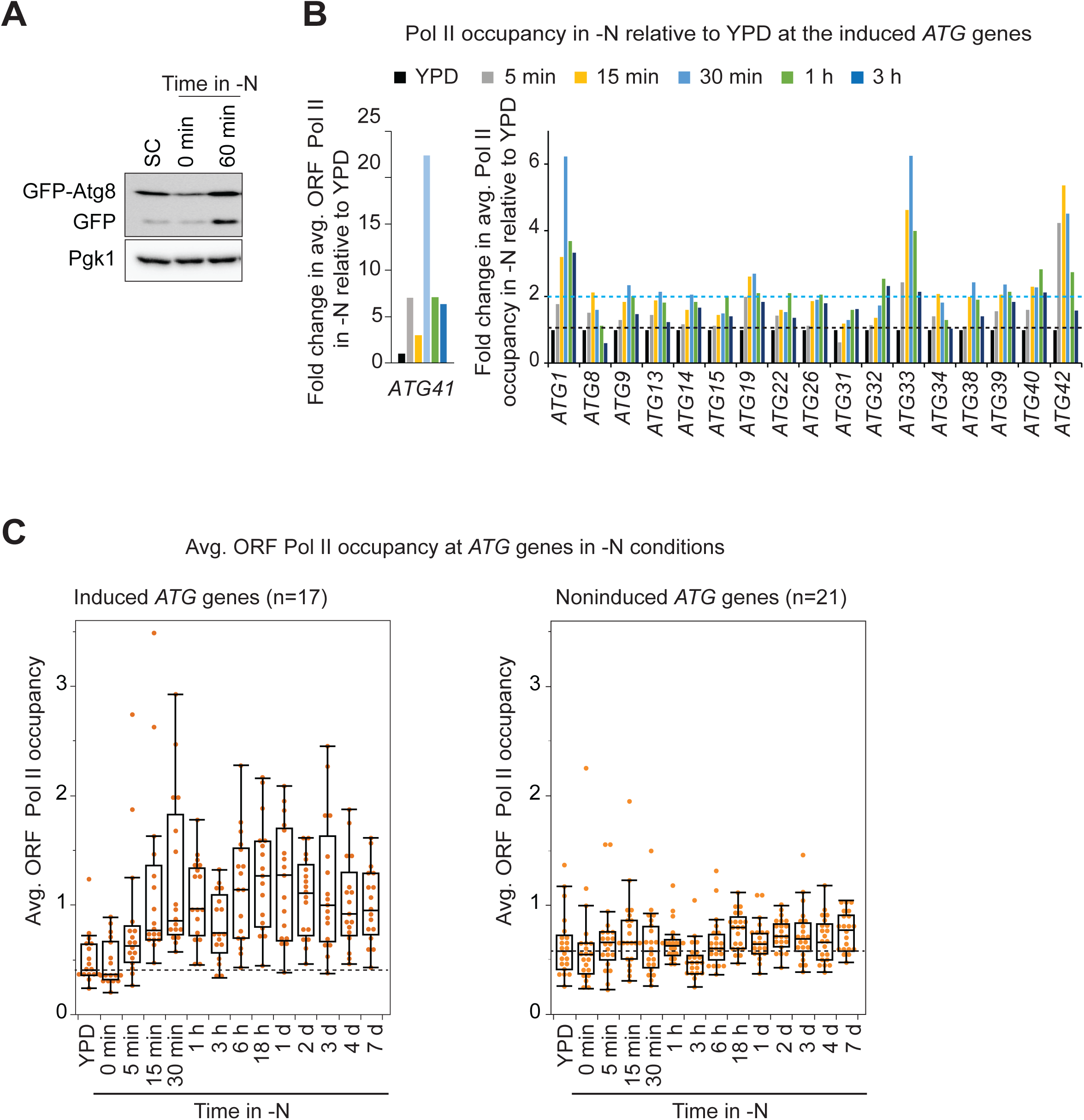
Several *ATG* genes are transcriptionally induced under N-starvation conditions. (**A**) WT cells harboring the GFP-*ATG8* plasmid were grown in synthetic complete (SC) media or subjected to -N media (0 and 60 min), and GFP signals were detected by western blot. Pgk1 is used as a loading control. (**B**) Fold changes in Pol II occupancy for *ATG41* (left panel) and other induced *ATG* genes (right panel) are shown. The fold change in Pol II occupancy for each induced *ATG* gene was calculated relative to the occupancy in YPD (set to 1). The genes shown exhibited at least a two-fold increase compared to YPD. (**C**) Average ORF Pol II occupancies in YPD and -N conditions (0-min to -d time points) are plotted for the induced *ATG* genes (left panel) and the non-induced *ATG* genes (right panel), excluding *ATG41*. Each dot represents average ORF Pol II occupancy for one *ATG* gene from two biological replicates.

Previous studies have shown upregulation of specific *ATG* genes in N starvation, including *ATG1*, *ATG8,* and *ATG41* [42–44]. However, our understanding of the activation of other *ATG* genes in -N conditions is limited. For instance, are all *ATG* genes upregulated, or are specific ones induced at a particular time during N starvation? To gain insight into this question, we examined Pol II occupancies at *ATG* genes. Notably, three *ATG* genes—*ATG8*, *ATG18*, and *ATG41*—ranked among the top 20% of the most transcribed genes in nutrient-rich medium, indicating their potential significance in basal autophagy in nutrient-rich environments (Fig. S2 and Table S3). We then analyzed Pol II occupancy at *ATG* genes across different starvation time points to identify the genes induced under these conditions (Fig. 2B and 2C). Based on induction levels, we classified *ATG* genes as induced *ATG* or noninduced *ATG*. The induced *ATG* genes group contained eighteen genes exhibiting at least a 2-fold increase in Pol II occupancy, minimally at one time point during -N starvation for the first 3 h (Fig. 2B). Most of these genes were induced within 30 min of starvation, except *ATG15, ATG22, ATG32, and ATG40,* which appeared to be most induced at the 1-h time point. Despite slight variations, the induction pattern of the induced *ATG* genes was quite similar, as all displayed a gradual increase in Pol II occupancy, peaking at either the 30-min or 1-h mark before significantly declining at the 3-h time point (p=0.05). Tracking Pol II occupancies over an extended period revealed that the dip in transcription at 3 h was temporary, as Pol II occupancies at these genes increased by the 6-h mark and were maintained until 18 h, with a slight decline at the later time points until day 7 (Fig. 2C). On average, the level of these genes remained greater than 2-fold at all time points, from 6 h to 7 days, compared to their levels in YPD. These Pol II averages exclude *ATG41*, which showed more than a 22-fold induction at the 30-min time point, beyond which its Pol II occupancy was comparable to *ATG1* and *ATG33* (Fig. 2B). Noninduced *ATG* genes showed a smaller induction, although the pattern of Pol II changes for these genes was similar to that of the induced *ATG* genes (Fig. 2C). Thus, the data show that approximately half of the known *ATG* genes are transcriptionally induced, with very similar kinetics, and they remain transcriptionally active for extended starvation periods.

### Transient downregulation of ribosomal biosynthetic genes during nitrogen starvation

RP genes are downregulated during various types of stress [45, 46]. Ribosome biosynthesis requires both RP genes (137 genes) and ribosome biogenesis genes (RiBi; ∼200 genes) [47]. The RiBi (*FCY2* and *YEF3*) and RP genes (*RPL8* and *RPS12*) that we examined exemplify the general pattern of transcriptional changes during nitrogen starvation (Fig. 3A). The Pol II occupancies on average declined immediately after shifting to -N conditions (0 min), resulting in a reduction of 37%, and continued to decrease progressively, reaching the minimum mean occupancies at 30 min, with a decline of more than 85% (Fig. 3B-C, Table S4). Thus, RP gene transcription was substantially reduced within 30 min of starvation; however, Pol II occupancy subsequently increased, reaching the levels seen in YPD by 3 h for the time points analyzed. The occupancy declined at the 6-h time point, falling below levels observed under YPD conditions. However, it remained higher than the 30-min time point (the lowest levels in -N), and these levels were then maintained throughout the remaining starvation period (Fig. 3C). The Pol II occupancy profile of RiBi genes in -N resembled that of RP genes (Fig. 3D and 3E). Pol II levels at RiBi genes were, on average, lower than RP genes (3.2 vs. 1.06 average normalized ChIP-seq reads); however, as with RP genes, they showed a substantial (75%) reduction in Pol II occupancy, dropping from 1.06 in YPD to 0.3 at the 30-min mark (Fig. 3D and 3E, Table S5). These results, showing the reactivation of genes involved in ribosome biosynthesis, are surprising because the TOR pathway promotes transcription of RP genes, and nitrogen starvation inhibits TOR [48, 49]. Although genes involved in ribosome biosynthesis are rapidly downregulated at the transcriptional level, the data indicate that these genes are reinduced and remain transcriptionally active during the extended starvation periods. It remains unclear which regulatory mechanisms trigger the reinduction of genes involved in ribosome biogenesis, or whether new ribosomes are produced, particularly under conditions of prolonged nitrogen starvation.

**Figure 3.**
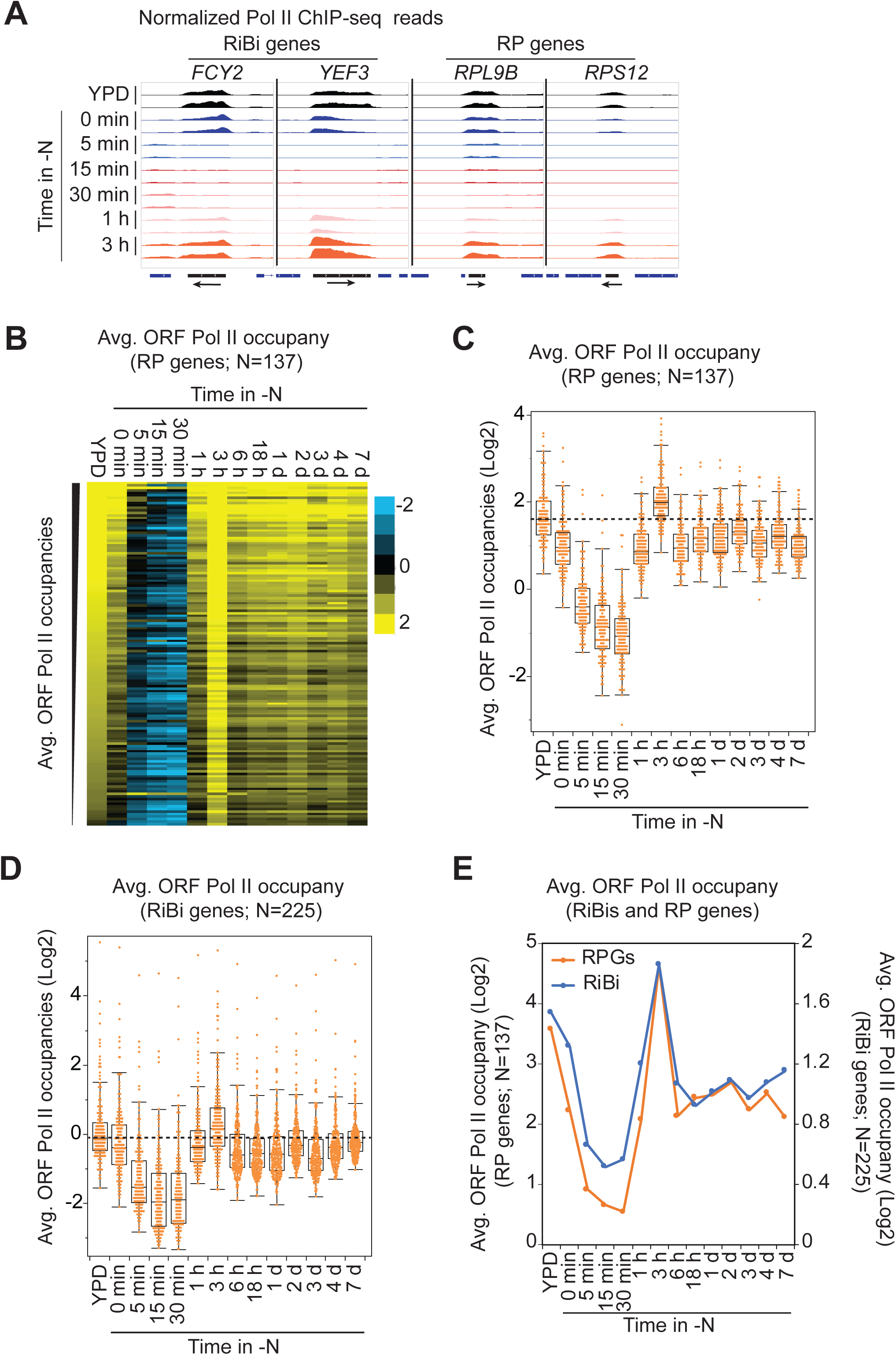
Ribosomal biosynthetic genes are transiently downregulated in nitrogen starvation conditions. (**A**) Genome browser screenshot showing normalized Pol II ChIP-seq reads for two representative ribosomal biogenesis (RiBi) genes and two ribosomal protein (RP) genes in YPD and from 0 min to 3 h under -N conditions. An arrow at the bottom of the panel depicts the direction of transcription for each gene, and a vertical dashed line separates the non-contiguous genes. (**B-C**) The average ORF Pol II occupancies in RP genes are displayed as a heatmap (B). The genes are sorted by their Pol II levels in the ORFs, from highest to lowest. The occupancies for RP genes are plotted in a boxplot (C). The dashed line represents the median Pol II occupancy for RP genes in YPD, and the individual orange dot represents the average ORF Pol II occupancy for one RP gene from two biological replicates. (**D**) The average Pol II occupancies for the ribosome biogenesis (RiBi) genes are represented as a boxplot. The dashed line represents the median Pol II occupancy in YPD, and the individual orange dot represents the average ORF Pol II occupancy for one RiBi gene from two biological replicates. (**E**) The similarity in induction of RP and RiBi genes is shown as a line plot tracking average ORF Pol II occupancies for all RP and RiBi genes over 7 days in -N conditions.

### Gcn4 target genes are upregulated in nitrogen-starvation conditions

While starvation induces autophagy, facilitating the recycling of amino acids, it also induces expression of Gcn4, a transcription factor for amino acid biosynthetic genes (Fig. 4A) [50–53]. Gcn4 also stimulates the transcription of *ATG1* and *ATG41* [43, 54]. Treatment of cells with sulfometuron methyl (SM), an inhibitor of valine and isoleucine biosynthesis, increases Gcn4 levels [50, 55–57] and induces transcription of its target genes, including most of the amino acid biosynthetic genes [51, 58, 59]. To determine which Gcn4 targets are induced during nitrogen starvation, we compared the genes induced by N starvation to those induced by SM using data from a previous study [59]. We identified 154 genes that showed more than a 2-fold increase in Pol II occupancies in both N-starved (30-min time point) and SM-treated cells (Fig. 4B). A significant proportion of these genes were involved in amino acid biosynthesis and metabolism (Fig. 4C-4D). The maximal induction for these genes was evident at 30 min, with an average induction of 4.9-fold (Fig. 4E). These results indicate that, alongside the activation of *ATG* genes, genes related to amino acid biosynthesis were upregulated during nitrogen starvation. Notably, this response was swift, with a nearly 5-fold increase in these genes within just 30 min (Fig. 4E). Among the most induced genes were those involved in arginine, methionine, asparagine, and leucine biosynthesis, as well as some of the *ATG* genes, such as *ATG1*, *ATG33*, and *ATG41*. While these genes exhibited early activation, 91 of the 154 genes continued to be expressed (showing over a 2-fold Pol II occupancy compared to YPD) even on day 7 of nitrogen starvation. This genome-wide analysis suggests that amino acid biosynthetic genes continued to be expressed alongside some *ATG* genes under prolonged starvation conditions and may play a role in amino acid biosynthesis under nitrogen-limited conditions.

**Figure 4.**
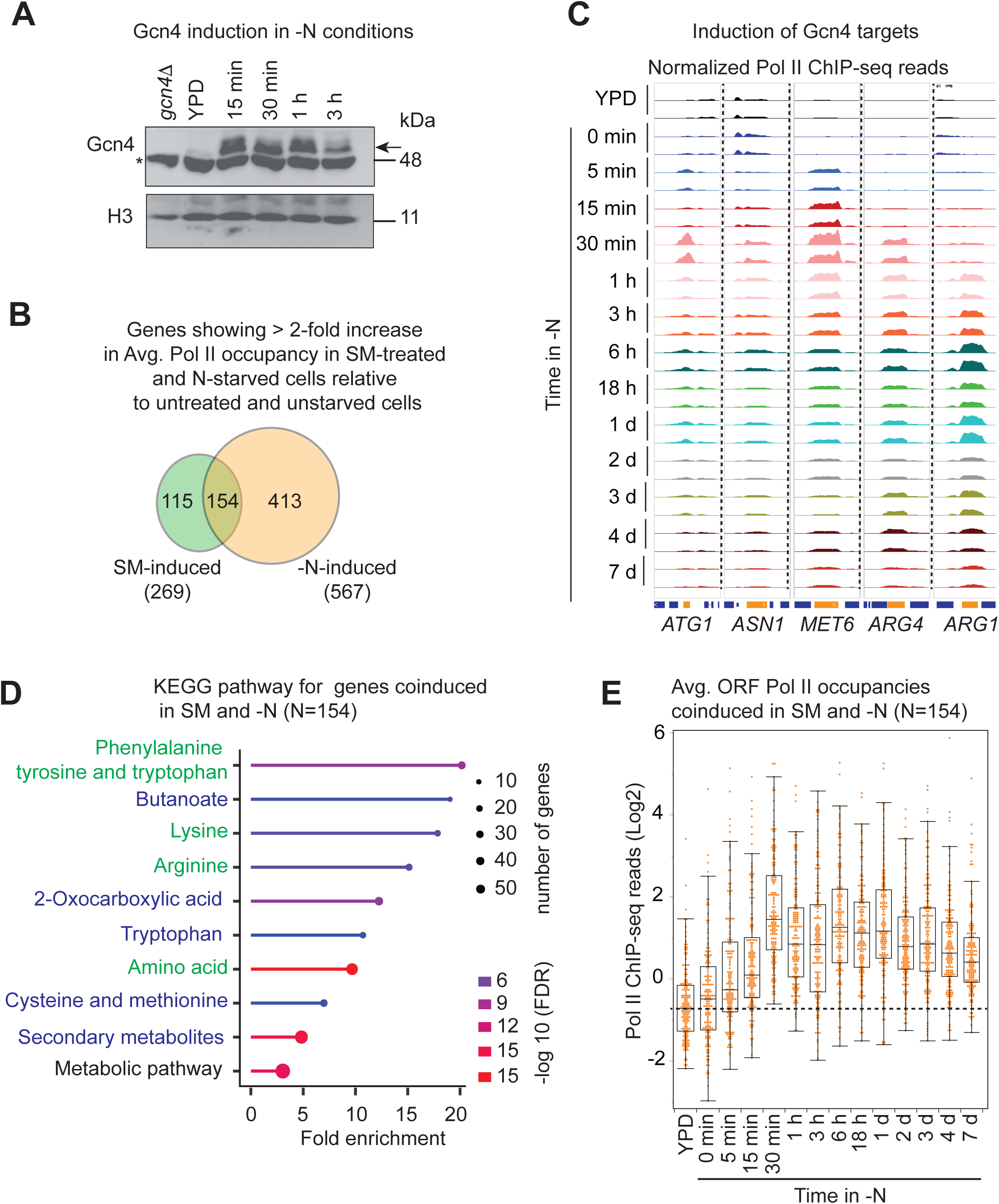
Many Gcn4 target genes, including amino acid biosynthetic genes, are induced under nitrogen-starvation conditions. (**A**) Western blot showing Gcn4 protein levels in nitrogen-starvation conditions. A *gcn4*Δ strain is used as a negative control, and histone H3 is used as the loading control. The asterisk represents a non-specific band detected by anti-Gcn4 antibodies. (**B**) Venn diagram showing genes induced by ≥2 fold in cells treated with the isoleucine/valine (ILV) inhibitor sulfometuron methyl (SM) and genes induced during nitrogen starvation. (**C**) Genome browser screenshot showing normalized Pol II ChIP-seq reads at five representative Gcn4-target genes in YPD and -N conditions from 0-min to 7-d time points. A vertical dashed line separates the non-contiguous genes. (**D**) Graphical representation of gene ontology analysis reveals that genes co-induced by SM and nitrogen starvation are enriched for amino acid biosynthetic pathways. (**E**) Pol II occupancies at the co-induced genes are shown as a box plot. The dashed line represents the median Pol II occupancy in YPD. Individual dots represent the average ORF Pol II occupancy for one Gcn4 target gene from two biological replicates.

### Nitrogen starvation conditions elicit global changes in histone occupancy

Yeast genes feature a nucleosome-depleted region (NDR) located upstream of the ORF, flanked by a minus one (-1; upstream) and a plus one (+1; downstream) nucleosome (Nuc) (Fig. 5A). During transcriptional induction, histones are evicted or repositioned from gene promoters to expose DNA to the transcriptional machinery [22, 57, 60], whereas histone occupancies increase upon a reduction in transcription [57, 61–63]. To analyze changes in chromatin structure in response to nitrogen starvation, we determined genome-wide histone H3 occupancy using ChIP-seq. The genome browser shots for H3 occupancies are shown for five representative downregulated genes in -N (Fig. 5B). All five genes showed increases in H3 occupancy in their promoter and 5’ ORF regions in -N conditions. The genome browser track showing the overlap of H3 occupancies in YPD and 30 min in -N reveals an increase in occupancy in the promoter regions of each gene (Fig. 5B, region pointed to by blue arrows). *HPT1* and *TDH2* additionally revealed substantial increases in H3 over their coding regions. To better evaluate histone occupancy changes relative to transcriptional changes, we computed H3 occupancies at the +1_Nuc and -1_Nuc positions and analyzed H3 occupancies for genes exhibiting at least a 2-fold reduction in Pol II occupancy at the 30-min time point during N starvation. The H3 occupancies at the +1_Nuc region showed a gradual increase over time in -N across all five genes, with smaller increases in the *FAA4* gene (Fig. 5C). Conversely, Pol II occupancies in their coding regions demonstrated a steady decrease over time, linking increased histone occupancies to transcriptional downregulation observed under N-starvation conditions.

**Figure 5.**
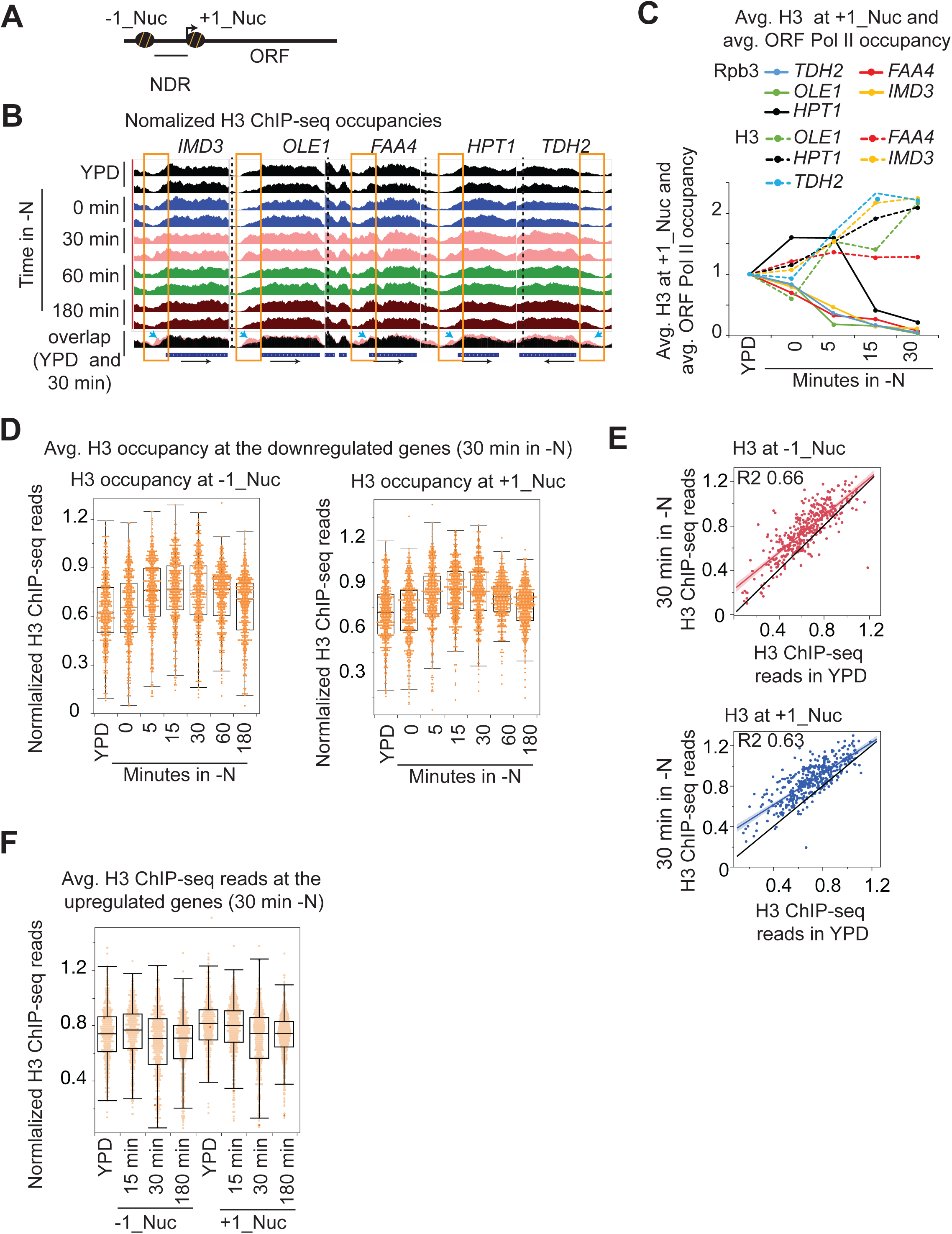
Global changes in histone occupancies under nitrogen starvation conditions. (**A**) Schematic showing the nucleosome-depleted region (NDR) upstream of the open reading frame (ORF). The nucleosome downstream of the NDR is known as the plus one (+1_Nuc), and the nucleosome upstream is minus one (-1_Nuc). (**B**) Genome browser shot showing normalized H3 ChIP-seq reads at five representative genes at various time points in -N, with the H3 occupancies around the NDR for each gene outlined by a rectangle. The bottommost track shows overlapping H3 ChIP-seq reads in YPD and at the 30-min time point in -N medium, with changes in H3 occupancy indicated by arrows. (**C**) The graph shows the average ORF Pol II (solid lines) and H3 ChIP-seq reads (dashed lines) from two biological repeats at the indicated time points in -N conditions for the genes displayed in panel A. (**D**) Box plot showing average H3 ChIP-seq reads at the -1_Nuc (left panel) and +1_Nuc (right panel) positions for the genes that show at least two-fold downregulation in Pol II occupancies at the 30-min time point in nitrogen starvation conditions. (**E**) Scatter plot showing that the genes that are downregulated by more than 2-fold show increased H3 occupancy at both -1_Nuc (top panel) and +1_Nuc positions (bottom panel). The black line shows no change. (**F**) Box plot showing average H3 ChIP-seq reads at the -1_Nuc and +1_Nuc positions for the genes that are upregulated in -N conditions at the 30-min time point. The individual dots represent the average H3 occupancy of one gene at the indicated nucleosome position from two biological replicates.

The downregulated genes showed a steady increase in H3 occupancy at both -1_Nuc and +1_Nuc positions during starvation, peaking at 15 to 30 min (Fig. 5D and 5E; and Tables S6 and S7). A reduction in H3 occupancy at the 3-h mark aligned with our finding of increased transcriptional induction of AA and ribosome biogenesis genes at that time. The increase in H3 at 30 min extended to most of the genes, although those with the lowest H3 occupancy in YPD conditions (most highly transcribed) exhibited a greater increase (Fig. 5E). In contrast to the downregulated genes, the induced genes (more than two-fold increase in Pol II occupancy at the 30-min time point in starvation) showed lower H3 occupancies at the -1 and +1 regions at the 30-min time point compared to YPD suggesting histone eviction occurred during the induction of these genes in -N medium. The changes in histone occupancy suggest that histone modifiers, histone chaperones, and chromatin remodelers may play a crucial role in eliciting transcriptional changes during nitrogen starvation and could also promote autophagy.

### Histone occupancy does not change at most ATG genes

Because many *ATG* genes were transcriptionally induced, we examined whether they also exhibited histone eviction during nitrogen starvation. We did not detect any significant changes in average H3 occupancy during N-starvation compared to nutrient-rich conditions for either the induced or non-induced *ATG* genes (Fig. S3A and S3B). However, H3 occupancy was substantially reduced at *ATG41*, and a smaller reduction was seen in the *ATG1* 5’ upstream region. However, *ATG8* and *ATG33* did not display such reductions (Fig. 6A) despite showing transcriptional induction during starvation (Fig. 2B). It is possible that the induction of some of the *ATG* genes involves nucleosome repositioning and not eviction, or that ChIP-seq fails to capture subtle changes in H3 occupancies that may accompany *ATG* gene induction. Nonetheless, two induced *ATG* genes showed reduced H3 occupancy in -N conditions.

**Figure 6.**
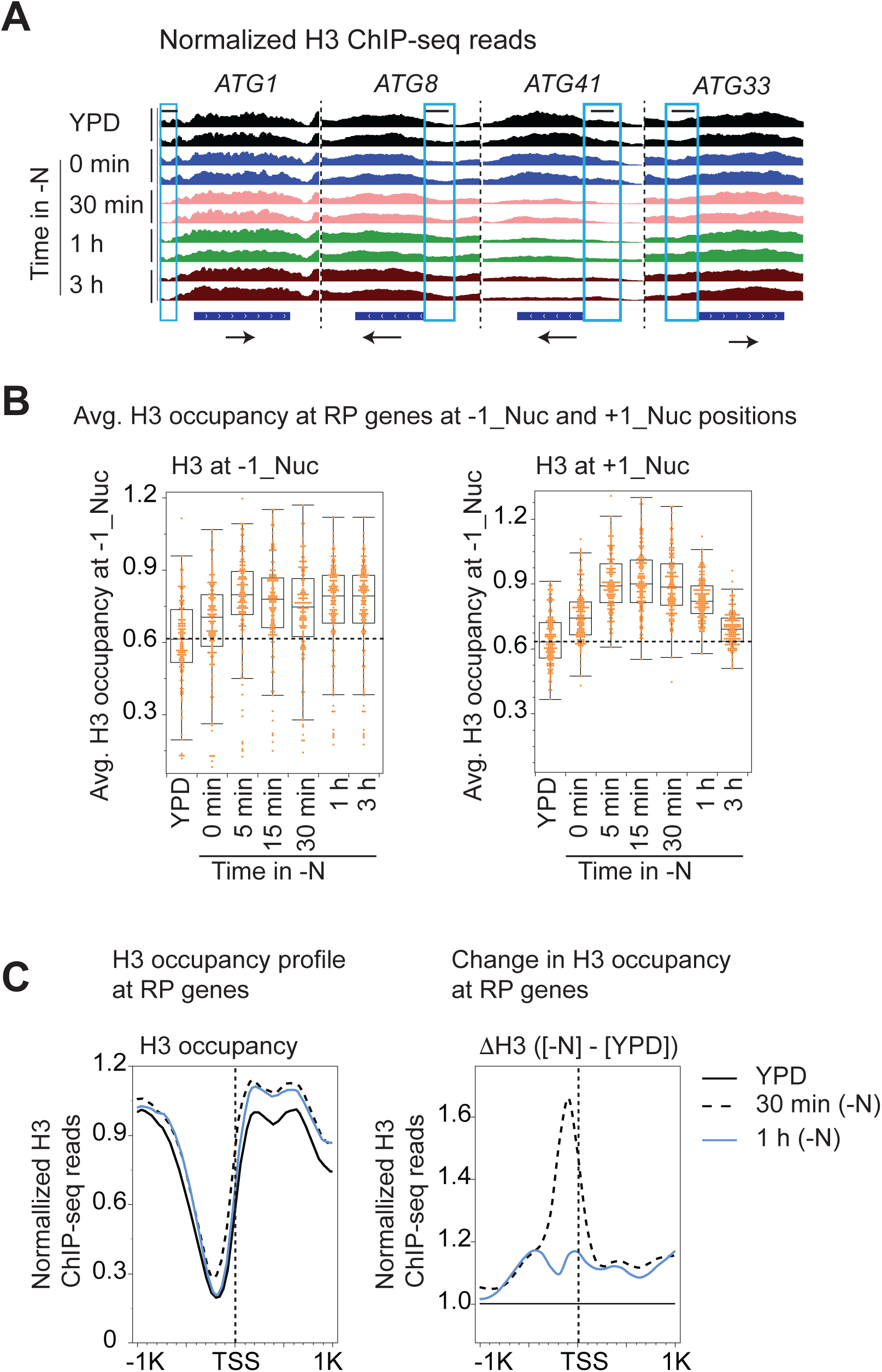
Ribosomal genes show an initial increase in histone occupancy followed by a reduction in their promoters. (**A**) Genome browser shot showing normalized H3 ChIP-seq reads during -N conditions at *ATG1*, *ATG8*, *ATG33,* and *ATG41*. The arrow depicts the direction of transcription, and the blue rectangle specifies the 5’ ORF and promoter region. The upstream region of the *ATG1* ORF showing the greatest reduction in H3 is marked by a black line. (**B**) Box plot showing average H3 ChIP-seq reads for each RP gene at specified positions (-1_Nuc or +1_Nuc) in YPD and nitrogen-starvation conditions. The individual dots represent the average H3 occupancy of a single gene at the specified position across two biological replicates. **(C)** Metagene plots showing H3 ChIP-seq reads 1 kb upstream and downstream of the transcription start sites (TSS) for RP genes (left panel). The differences in H3 occupancies (ΔH3; [H3 in -N] – [H3 in YPD]) are plotted in the right panel.

### Reinduction of ribosomal biosynthetic genes during nitrogen starvation involves histone eviction from their promoters

RP genes exhibit transcriptional downregulation within 30 min, followed by upregulation at later time points (Fig. 3B and 3C). Consequently, we investigated whether the rapid downregulation and subsequent activation are associated with changes in histone occupancy. The H3 occupancies at the -1_Nuc and +1_Nuc for RP genes were significantly increased as early as the 0-min time point and continued to increase until the 15-min time point (Fig. 6B). The rapid increase in H3 occupancies aligned with the substantial reductions in Pol II occupancies at RP genes in -N conditions. Furthermore, consistent with the reinduction of RP genes, H3 occupancy began to decline after 30 min, reaching the levels observed in YPD medium at the +1_Nuc position by the 3-h time point in N-starvation (Fig. 6B; right panel). However, we did not see any significant changes in the H3 occupancy at the -1_Nuc position at the 3-h time point.

Because the +1_Nuc flanks the NDR, we asked whether the repression of RP genes involves filling in the NDR. The H3 occupancy profiles revealed that the NDR became narrower and shallower within 30 min of starvation. This could result from the movement of nucleosomes inwards towards the NDR (narrower), the assembly of nucleosomes with the NDRs (shallower), or a combination of both. However, by the 1-h time point, the profile resembled that of the cells in YPD (Fig. 6C), consistent with the reinduction of RP genes. The calculated differences in H3 occupancies (ΔH3; [H3 occupancies in -N conditions] – [H3 occupancy in YPD]) at each base pair at the 30-min and 1-h time points revealed that the maximal H3 increases were localized upstream of the TSS at the 30-min time point (Fig. 6C, right panel). The higher H3 occupancy probably impedes the assembly of the transcription preinitiation complex, leading to the downregulation seen at 30 minutes. However, H3 levels were significantly reduced at the 1-h time point, coinciding with increased transcription of RP genes. Thus, the data suggest that both the repression of RP genes in starvation and their subsequent induction involve substantial changes in histone occupancies at their promoters.

### Altered histone acetylation to drive transcriptional reprogramming during nitrogen starvation

Acetylation of the histone N-terminal tails weakens the electrostatic attraction to DNA, and acetylated histones also help in recruiting bromodomain-containing ATP-dependent chromatin remodelers to reposition or remove histones [57, 64, 65]. Because transcriptional induction under N-starvation conditions was accompanied by histone eviction, we sought to determine the extent to which histones at induced genes were acetylated. Because Esa1 promotes autophagy and acetylates H4, we determined H4Ac levels by ChIP-seq in YPD or -N conditions (Table S8). We observed increased H3 occupancies at the downregulated genes and reduced H3 occupancies at the induced genes (Fig. 5 and Fig. 6). Therefore, to account for altered histone occupancies, we normalized H4Ac levels by H3 occupancy. We observed reductions in total H4Ac and H3-normalized H4Ac (H4Ac:H3) levels at +1_Nuc across the genome (Fig. S4A). Thus, both transcription and H4Ac declined globally in the -N condition. However, many genes that were poorly transcribed in YPD exhibited an increase in H4Ac near their transcription start site (TSS) in -N conditions (Fig. S4B; compare the top and bottom halves of heatmaps, which are arranged based on decreasing Pol II occupancy). The latter results might suggest that induced genes are acetylated, and the data support the idea that chromatin reorganizes to adapt to N-starvation conditions.

We next examined H4Ac at the induced RP, AA, and *ATG* genes. Considering that RP and AA genes showed changes in histone occupancy, we plotted H4Ac:H3 around their TSS. RP genes demonstrated a significant increase in H4Ac:H3 immediately downstream of the TSS at the +1_Nuc position. Notably, the +1_Nuc region showed nearly 2.5-fold higher H4Ac:H3 at the 3-h time point compared to the cells in YPD (Fig. 7A). It is worth noting that the total H3 levels at the 3-h time point were slightly higher than in YPD conditions (Fig. 6C). These observations suggest that the +1_Nucs at RP genes were hyperacetylated relative to those in nutrient-rich conditions and may stimulate the induction of RP genes in nitrogen-limited conditions. The Gcn4 targets (co-induced in -N and SM; Fig. 4B) also showed higher H4Ac:H3 ratios (Fig. 7B), indicating that histones at promoters were acetylated and evicted. Thus, acetylation of H4 is positively correlated with induction and histone eviction.

**Figure 7.**
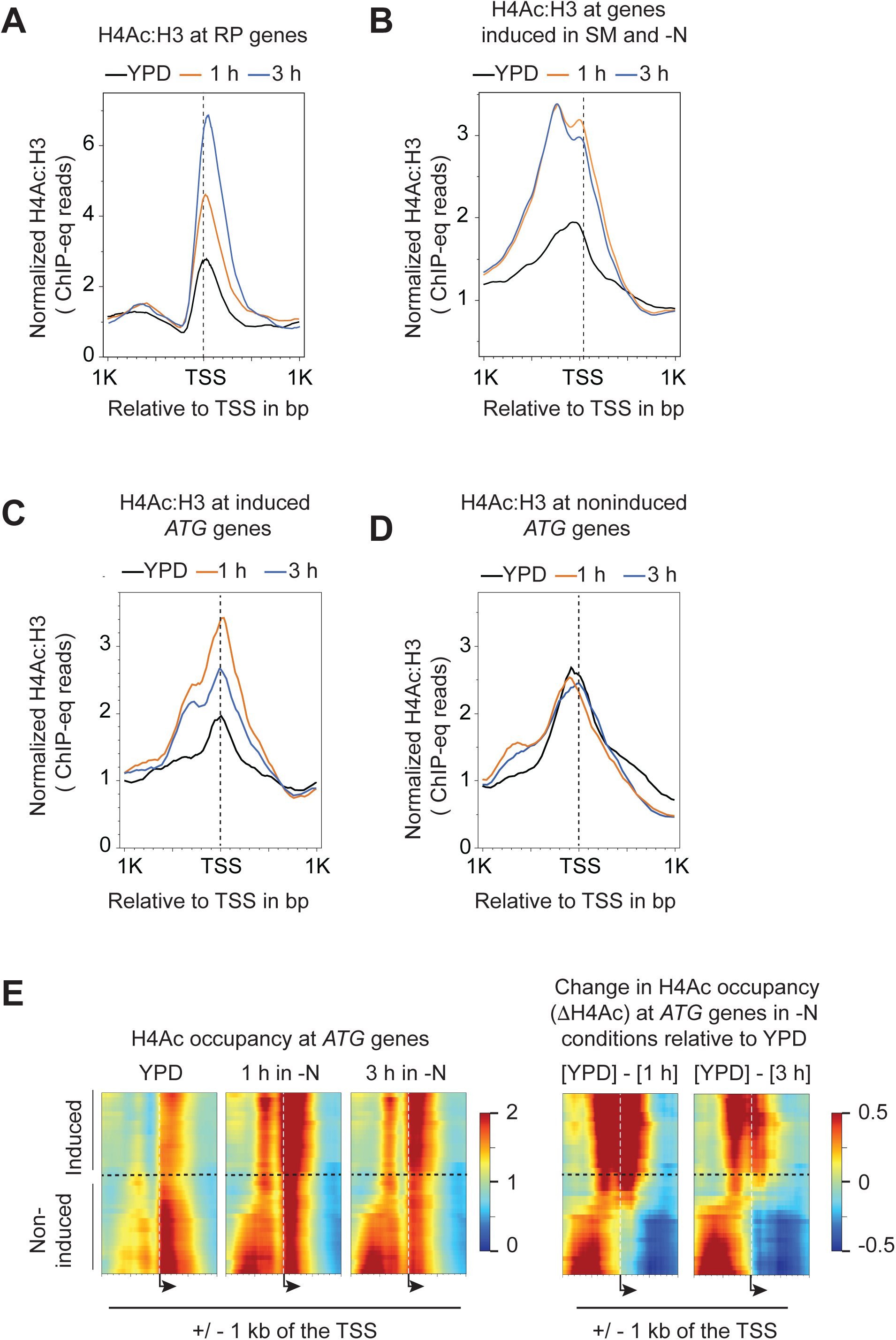
Histone H4 is acetylated during the induction of RP genes, Gcn4-targets, and *ATG* genes during N starvation. (**A-B**) Metagene plots showing acetylated H4 (H4Ac) ChIP-seq reads normalized to H3 ChIP-seq reads (H4Ac:H3) at RP genes (A) and for the SM and -N co-induced genes in YPD, and in -N media (1 h and 3 h) (B). The H4Ac:H3 levels are plotted +/- 1 kb around the TSS. (**C-D**) H4Ac:H3 levels in YPD and - N (1 h and 3 h) are plotted for the induced *ATG* genes (left panel) and non-induced *ATG* genes (right panel). (**E**) Heatmaps showing total H4Ac +/- 1 kb upstream and downstream of the TSS for *ATG* genes (left panels). The changes in H4Ac (ΔH4Ac, [H4Ac in -N] – [H4Ac in YPD]) are depicted as heatmaps. A dashed line separates induced and noninduced *ATG* genes in the heatmaps.

We finally examined whether *ATG* genes also show changes in H4Ac. These genes exhibited only minor changes in H3 occupancy (Fig. 6 and S6). Among these, the induced *ATG* genes exhibited an increase in H4Ac:H3 during starvation, with the highest levels observed at the 1-h time point. In contrast, consistent with the largely uniform Pol II occupancies in YPD and -N conditions, noninduced *ATG* genes did not reveal substantial changes in H4Ac:H3 in starvation conditions (Fig. 7E). The heatmaps depicting H4Ac revealed that the noninduced *ATG* genes harbored higher levels of acetylated H4 both at the -1_Nuc and +1_Nuc positions in YPD conditions compared to the induced *ATG* genes (Figure 7E; compare the top half with the bottom half of each heatmap). The latter genes exhibited a significant increase in H4Ac at both -1_Nuc and +1_Nuc at the 1-h and 3h time points. As such, our data show that induction of RP genes, AA genes, and *ATG* genes during N starvation is accompanied by increased histone H4 acetylation.

### Esa1 is critical for transcriptional induction in nitrogen-starvation conditions

Having observed the increased H4Ac at the induced genes, we sought to assess the role of Esa1 in this process. To achieve this, we generated an Esa1 auxin-inducible degron (AID) strain. Adding indole-3-acetic acid (IAA) for 60 min led to substantial depletion of Esa1 in YPD conditions (Fig. 8A). We then measured Pol II ChIP-seq occupancies in untreated and IAA-treated (*esa1*) cells in both YPD and -N media for 1 and 3 h.

**Figure 8.**
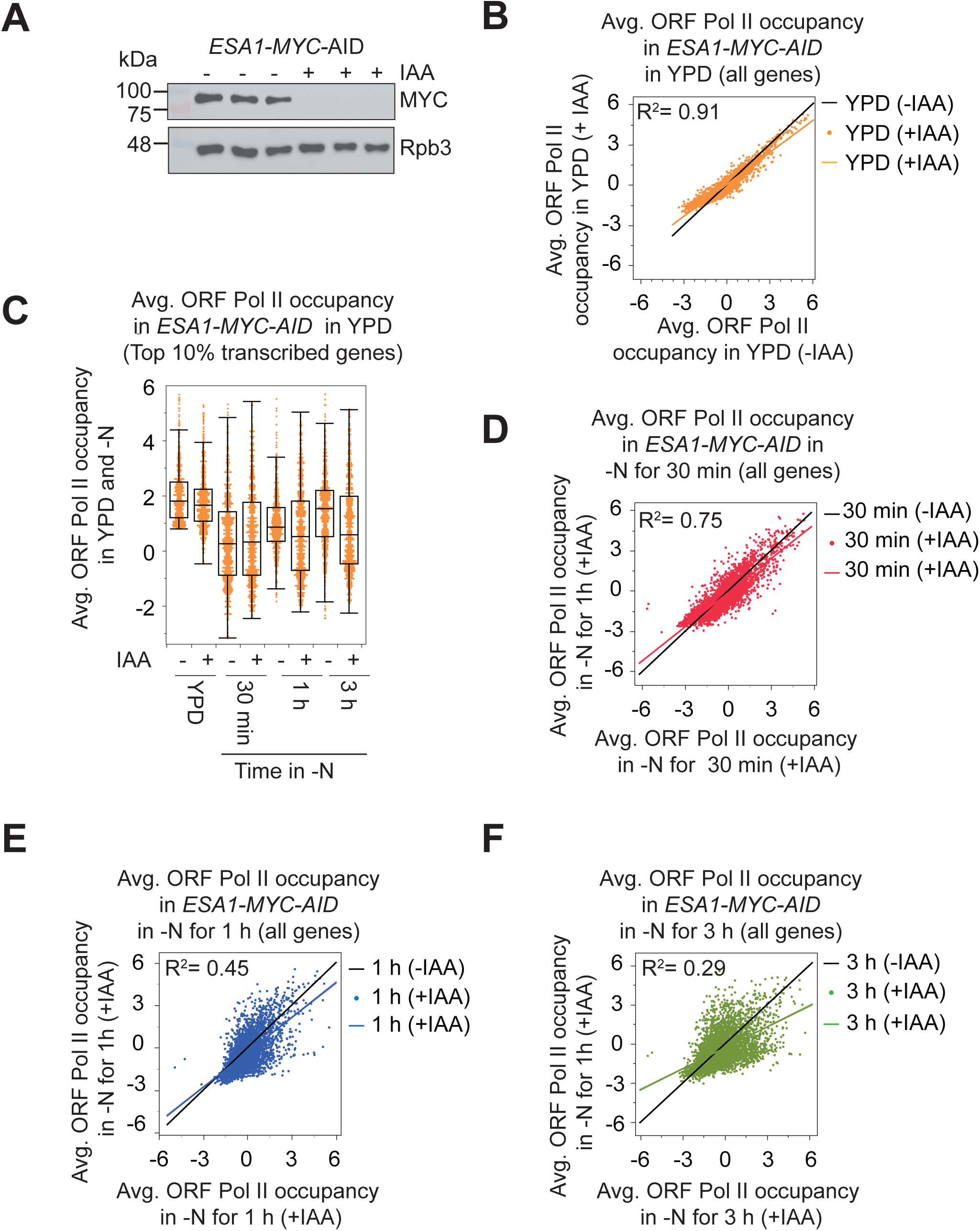
Esa1 promotes transcription during nitrogen starvation. (**A**) A Western blot showing substantial depletion of Esa1 in the *ESA1-MYC-AID* strain across three biological replicates in 60 min after treatment with 3-indole acetic acid (IAA). (**B-E**) Average ORF Pol II ChIP-seq reads for yeast genes (N=5620) in Esa1-depleted cells (+IAA) are compared to the reads in untreated cells in YPD (B) and nitrogen starvation for 30 min (C), 1 h (D), and 3 h (E). The coefficient of determination (R^2^) is provided for each comparison. **(F)** The average ORF Pol II ChIP-seq reads (Log2) for the top 10% of Pol II-occupied genes are displayed as box plots for -IAA and +IAA treated cells grown in YPD, or nitrogen-starved for 30 min, 1 h, and 3 h.

We first examined the impact of *esa1* on genome wide Pol II occupancies in YPD (Table S9). A small but significant change in Pol II occupancies (p = 7.8E-11) was observed in the *esa1* cells compared to untreated cells (R² = 0.91; Fig. 8B). This suggests that Esa1 has only a minor impact on transcription under nutrient-rich conditions. This finding was also true for the top 10% transcribed genes. Following Esa1 depletion, there was a small reduction in average Pol II occupancy for these genes, with the mean normalized ChIP-seq reads dropping from 1.62 to 1.43 (p=2E-4) (Fig. 8C; YPD). Of these, just 25 genes exhibited over a 2-fold reduction, while 81 genes showed more than a 1.5-fold decrease in Pol II occupancy, indicating that Esa1 promotes transcription of a small number of genes in nutrient-rich conditions.

We then examined the effect of Esa1 depletion on transcription in -N. Because Pol II occupancy decreased globally when cells were starved, we compared their occupancies at the same time points in -N conditions with and without IAA. The Pol II occupancy correlations between IAA-treated and untreated cells decreased gradually with increasing time in -N: 30 min; R²=0.75, p=1.9E-3 (Fig. 8D); 1 h; R²=0.45, p=8E-59 (Fig. 8E); and 3 h; R²=0.29, p=1.5E-9 (Fig. 8F). The significant reductions in Pol II occupancy in *esa1* cells underscored a crucial role for Esa1 in maintaining a normal transcriptional profile under starvation conditions. As noted above, no significant changes in Pol II occupancy were observed at 30 min for the top 10% transcribed genes (Fig. 8C). Because transcription of most genes is downregulated at 30 min, this observation may suggest that Esa1 plays no significant role in transcriptional downregulation under starvation conditions. By contrast, the Pol II occupancies at these genes were significantly lower in *esa1* cells at the 1-h (p=6.3E-5) and 3-h (p=8.1E-12) time points, suggesting that Esa1 is crucial for transcription of genes upregulated in -N conditions.

### Esa1 is required for the reinduction of ribosomal biosynthesis genes

We next examined the effect of Esa1 depletion on Pol II occupancy at genes induced under -N conditions. Genome browser snapshots of Pol II occupancy at *ARG1*, *ATG41*, *RPL3*, and *RPL9B* are shown (Fig. 9A). Pol II occupancies were significantly reduced in the IAA-treated cells at Gcn4 targets *ARG1* and *ATG41* as well as at RP genes *RPL3* and *RPL9B* (Fig. 9A; compare -IAA and + IAA in -N). Analysis of all RP genes in YPD revealed a 17.2% reduction (p < 0.006) in average Pol II occupancy in *esa1* cells (Fig. 9B), suggesting that Esa1 made a small but significant contribution to the transcription of RP genes in nutrient-rich conditions. In -N conditions, the Pol II at RP genes declined to a very similar extent in Esa1-containing and -depleted cells (Fig. 9B). However, RP genes failed to be transcriptionally induced in *esa1* cells as evident by markedly reduced levels of Pol II at these genes at the 1-h and 3-h time points (Fig. 9B). Compared to the 30-min time point, Esa1-containing cells showed a 4.1-fold increase in Pol II occupancy at 1-h and a 9.0-fold increase at the 3-h time point. By contrast, in *esa1* cells, RP genes were induced by 1.2- and 1.5-fold at the same time points. Likewise, the RiBi genes also failed to be induced in *esa1* cells, showing only a 1.1- and 1.2-fold increase at the 1-h and 3-h time points, respectively, compared to a 2.8- and 4.0-fold increase in nondepleted cells (Fig. 9C). Overall, the data indicate that Esa1 plays a critical role in reactivating RP genes and RiBi genes under starvation conditions.

**Figure 9.**
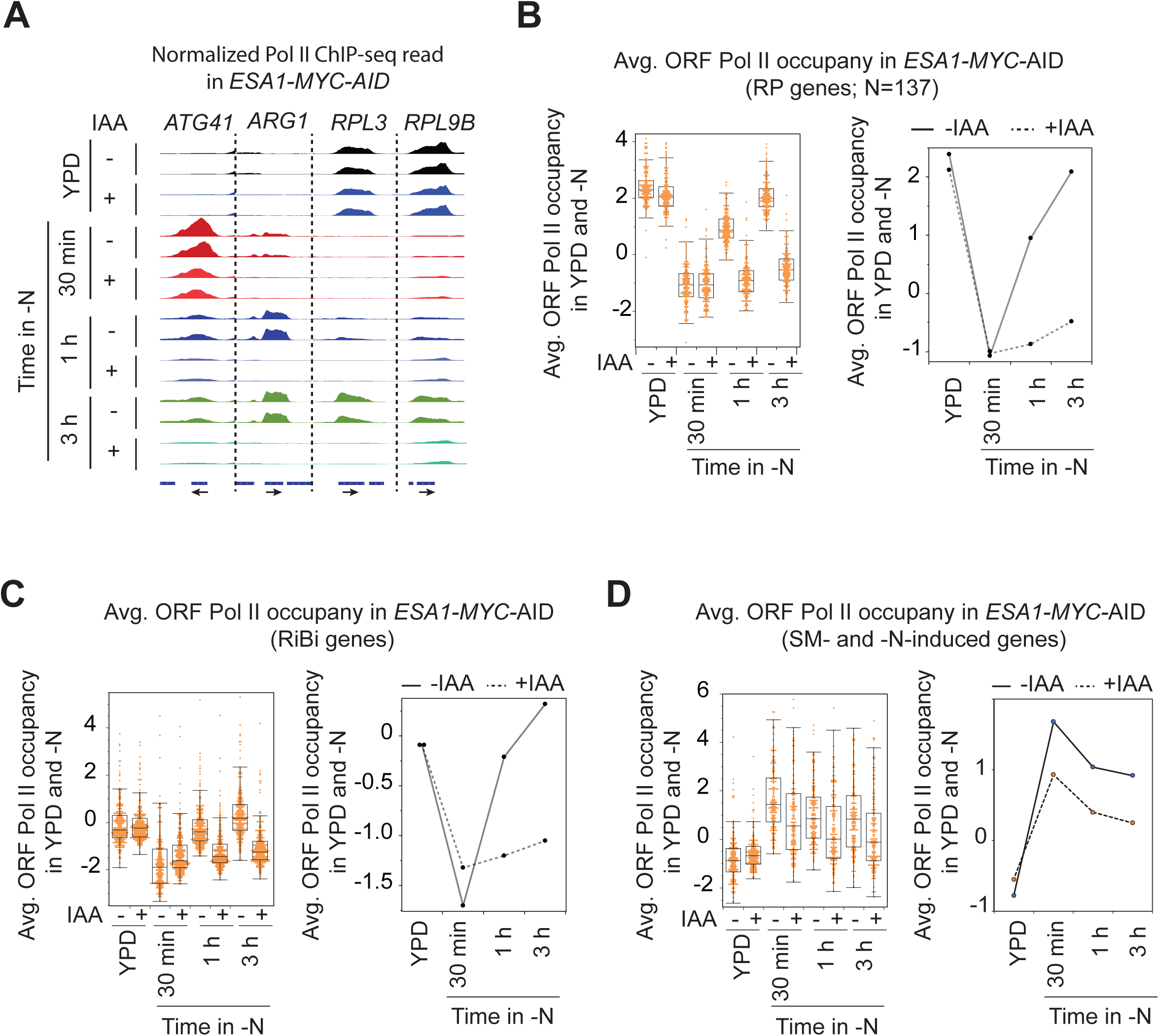
Esa1 plays a crucial role in the induction of ribosomal and Gcn4-target genes during nitrogen starvation. (**A**) Genome browser shot showing normalized Pol II ChIP-seq reads at representative genes in *ESA1-MYC-AID* cells (+/- IAA) grown in YPD and nitrogen-starved for 30 min, 1 h, and 3 h. (**B-D**) The average ORF Pol II ChIP-seq reads (Log2) in the *ESA1-MYC-AID* cells (+/- IAA) are shown as a boxplot (left panel) or line graph (right panels) for RP genes (B), RiBi genes (C), and SM and -N co-induced genes (D).

Many Gcn4 target genes are also activated under the -N condition. Therefore, we examined whether Esa1 deficiency reduced transcription of these genes. Esa1 depletion resulted in a 1.9 to 1.8-fold lower induction of AA biosynthetic genes from 30 min to 3-h under starvation conditions (Fig. 9D). Depleting Esa1 had no significant impact in YPD medium because these genes only showed basal expression under nutrient-rich conditions. Furthermore, given that these genes showed increased H4 acetylation (Fig. 7C), the reduced induction may suggest that Esa1-dependent H4 acetylation at these promoters was necessary for full induction of these genes, particularly during N starvation.

## Discussion

In this study, we investigated transcriptional changes and chromatin modifications induced by nitrogen starvation. Our results reveal that most genes are rapidly downregulated in starvation conditions, while a specific subset, including *ATG* genes, is upregulated. Although genes related to ribosomal proteins and biogenesis are initially downregulated, they are reactivated even as cells remain in starvation. While nitrogen starvation triggers autophagy, we observed a substantial increase in the expression of genes involved in amino acid biosynthesis, which remained high throughout the extended starvation period. Starved cells exhibit widespread changes in histone occupancy during starvation, with downregulated genes showing increased histone occupancy at their promoters and induced genes showing reduced occupancy and increased H4 acetylation. Furthermore, this study reveals the role of Esa1 in promoting activation of both ribosomal and AA genes under starvation conditions.

### Increased histones and reduced acetylation are associated with transcriptional downregulation in nitrogen-starvation conditions

We noted a swift downregulation of genes as soon as the cells were transferred to starvation medium (Fig. 1). The genes with the most significant declines in Pol II occupancy also exhibited notable increases in histone occupancy at their promoters and at the +1_Nuc position (Fig. 5B-5E), indicating that rapid repression may be promoted by increasing nucleosome assembly at the promoters. The increased +1_Nuc occupancy is likely to interfere with preinitiation complex/PIC assembly and binding of Spt15/TATA binding protein, because the +1_Nuc position generally harbors a TATA box in *S. cerevisiae* [66]. We also observed reduced H4 acetylation at the promoters of genes expressed under nutrient-rich conditions and repressed in -N conditions. It is possible that HDACs, such as the Rpd3 or Hos2-Set3 complexes [22, 67], are recruited to deacetylate nucleosomes during nitrogen starvation. Because these factors recognize methylated H3 [67–69], it will be interesting to determine whether the repressed genes also show increased histone methylation or whether the recruitment of these complexes is mediated in a methylation-independent manner.

### Histone eviction and elevated H4 acetylation occur at the majority of induced genes during nitrogen-starvation conditions

The data also reveal that genes involved in ribosome biogenesis, amino acid biosynthesis, and autophagy were induced during nitrogen starvation. The vast majority of the induced genes showed reduced histone occupancy (Fig. 5F), indicating that histones are evicted during transcriptional induction. Additionally, these genes showed increased H4Ac, suggesting a possible coordination between histone acetylation and ATP-dependent chromatin remodelers to evict histones. A chromatin remodeling complex containing Ino80 (Ino80C) affects autophagy negatively [26]. Ino80 remodels H2A.Z-containing nucleosomes, which are found primarily at the +1_Nuc [70, 71]. However, Ino80C also evicts canonical H2A-containing nucleosomes under starvation conditions [61]. It remains to be seen whether Ino80 plays a role in evicting H2A from the promoters of genes induced during nitrogen starvation.

The H4 HAT, Esa1, promotes autophagy [27]. The data show that Esa1 is required for the induction of ribosomal genes (Fig. 9), and these genes exhibit higher H4Ac levels in nitrogen starvation than in nutrient-rich media. H4Ac could promote recruitment of bromodomain-containing chromatin remodeling complexes such as SWI/SNF or RSC. These complexes evict histones from the promoters of ribosomal and AA biosynthetic genes [63, 65, 72, 73]. In fact, eviction of histones from amino acid biosynthetic genes during AA starvation relies on SWI-SNF and RSC [51]. Because a subset of *ATG* genes showed increased H4Ac, it is tempting to speculate that RSC and SWI/SNF remodel chromatin in the *ATG* gene promoters, thereby facilitating their induction. However, the extent to which Esa1, SWI/SNF, and RSC cooperate to regulate *ATG* gene transcription remains to be investigated. Furthermore, the data suggest that Esa1 enhances the transcription of numerous genes during nitrogen starvation. It will be important to study how the activation of Esa1-dependent genes influences autophagy and cell survival in acute starvation conditions. These investigations could reveal a broader role for Esa1 beyond its established function in acetylating Atg3, suggesting that Esa1 may also facilitate autophagy by activating specific genes under nitrogen-starved conditions.

### Induction of ATG genes

Considering that nitrogen starvation strongly induces autophagy [6, 13], we anticipated a significant increase in *ATG* gene levels. Surprisingly, however, most *ATG* genes showed only a modest increase in Pol II occupancy compared with the strong induction observed in ribosomal and amino acid biosynthetic genes, except for one gene, *ATG41*. Our findings reveal a subset of *ATG* genes that are induced during nitrogen starvation with remarkably similar kinetics. *ATG* genes are regulated by a diverse set of transcription factors, including Gcn4, Gln3 and Gat1 [33, 43, 74, 75]. Hence, it is unclear how nitrogen starvation coordinates the transcriptional induction of *ATG* genes. We surmise chromatin remodelers and histone modifications may serve as key regulators for the coordinated expression of these genes. This conclusion is supported by data showing that the +1_Nuc of these genes exhibit increased H4Ac levels in SD-N.

### Amino acid and ribosome biosynthetic genes are induced under nitrogen starvation conditions

Cellular components are degraded via autophagy, and, thus, become vacuole-released precursor molecules that can be reused. Under starvation conditions, proteins are degraded to recycle AA that can be utilized for protein synthesis. Our data show that AA biosynthetic genes are strongly upregulated, suggesting that, in addition to autophagy [34], the cells may also upregulate de novo amino acid biosynthesis. Ribosomes are one of the most abundant protein complexes in cells, and it is thought that degradation of ribosomes during starvation could provide the necessary AAs for protein synthesis. Our results are consistent with previous studies showing that nitrogen-starved cells channel amino acids recycled via autophagy to sustain glutamate synthesis, and that certain amino acid syntheses, such as for serine, do not depend on autophagy [35]. Additionally, ribophagy, the selective degradation of ribosomes, does not help maintain the basic amino acid pool [76]. Therefore, the expression of amino acid biosynthetic enzymes helps replenish amino acid pools, even during active autophagy.

Ribosomal biosynthetic genes, particularly the structural RP genes, are downregulated in response to various stresses, such as amino acid starvation and heat shock [77–79]. However, we note that the decrease in transcription of ribosomal biosynthetic genes was only temporary in -N conditions. At the 3-h time point, the Pol II levels at the RP genes are similar to those in YPD. Because starvation suppresses the TOR pathway, the subsequent re-induction of genes suggests that TOR inhibition may be relieved to some extent to sustain ribosomal gene expression even under the N-starvation stress. TOR activity is influenced by amino acid levels [80, 81], and increased amino acid biosynthesis [82] through transcriptional activation of amino acid biosynthetic genes, coupled with the potential release of amino acids via autophagy-mediated protein degradation, could help in alleviating TOR inhibition under starvation conditions. Our research indicates that during nitrogen starvation, cells transcribe at least three categories of genes: (1) those related to autophagy, (2) those involved in amino acid biosynthesis, and (3) those associated with ribosome biosynthesis. This finding suggests that under these conditions cells not only prioritize activating catabolic pathways such as autophagy but also induce genes involved in anabolic processes.

## MATERIALS AND METHODS

### Yeast Strain Construction

All *Saccharomyces cerevisiae* strains and plasmids used in this study are listed in Table S1. *ESA1*-MYC-AID tagged cells were generated by a PCR-based method [83] using the plasmid pHis-AID-9myc (Addgene, 99524; deposited by Dr. Helle Ulrich’s lab) as described previously [57]. Briefly, primers containing homologous regions flanking the stop codon of the *ESA1* gene were used to amplify the AID-MYC tag, and cells containing a *TIR1* gene were transformed using the lithium acetate method [84]. *TIR1* was inserted into the *LEU2* locus after digesting pTIR1 with the Pme1 enzyme [85]. Successful transformants were verified by expression of the tagged protein and depletion upon addition of 3-indoleacetic acid (IAA; Millipore Sigma, I3750).

### Yeast Cell Growth

Cells were grown in YPD (1% yeast extract, Fisher, BP1422-2; 2% peptone, Thermo Scientific, J20048; 2% dextrose, Fisher Bioreagents, BP350) or synthetic complete (SC) media. The SC base was prepared using 0.2% amino acid mixture (prepared from 2 g of inositol, .5 g of adenine and .2 g of para-aminobenzoic acid and 2 g of L-amino acid mix prepared from 2 g of each amino acid [Millipore Sigma], except arginine, isoleucine/valine [ILV], leucine, methionine, tryptophan, and histidine), 1.54 g of yeast nitrogen base without amino acids (YNB w/o AA, Millipore Sigma, Y0626), 4.5 g of ammonium sulfate (Macron 3512-12), and 2 % dextrose. The remaining amino acids and uracil were added to the SC base (methionine 0.5 mM, leucine 1 mM, ILV 0.25 mM, tryptophan 0.2 mM, arginine 0.25 mM, uracil 0.25 mM, and histidine 0.12 mM). For SC - URA, uracil was omitted from the media. The cells were grown at 30°C to an absorbance A_600_ of 0.6-0.7 and collected for further analysis or washed with sterile deionized water and transferred to starvation medium lacking amino acids and nitrogen (SD-N; 2% dextrose [Fisher] and .17% of YNB without AA and nitrogen, Difco 233520). Conditional depletion of Esa1 was achieved by adding IAA (1 mM) to *ESA1*-AID-MYC cells 90 min before transferring them to -N medium. Cells were incubated in SD-N medium for several time points ranging from minutes to days at 30°C with shaking. At each selected time point, cultures were spiked-in using *Schizosaccharomyces pombe,* grown in YES medium (0.5% yeast extract, Fisher, BP1422-2; 3% dextrose [Fisher], and 0.02% of adenine, uracil, histidine, leucine, and lysine; Millipore Sigma) to a final A_600_ of 15% of the A_600_ of the *S. cerevisiae* cultures. Following spike-in, cultures were immediately crosslinked with a final concentration of 1% formaldehyde in the crosslinking buffer (50 mM HEPES-KOH, pH 7.5, 1 mM EDTA, and 100 mM NaCl) for 15 min, and crosslinking was quenched with 15 mL of 2.5 M glycine. Crosslinked cells were collected by centrifugation at 2000 x g for 5 min at 4°C, washed twice with chilled Tris-buffered saline (TBS; KD Medical, RGF-3346), and stored at -80°C until further use.

### Preparation of Chromatin from Crosslinked Cells

Crosslinked cells were resuspended in 500 µL of chilled FA lysis buffer (50 mM HEPES-KOH, pH 7.5, 1 mM EDTA, 140 mM NaCl, 1% Triton X-100 [Millipore Sigma, T9284], 0.1% sodium deoxycholate [Millipore Sigma, 30970] containing freshly added protease inhibitor cocktail (FA-PIC; 1 µg / ml of pepstatin A [Thermo Scientific, 78436], 1 µg / ml of leupeptin [Thermo Scientific, 78435], and 1 µg / ml of pepstatin A [Thermo Scientific, 78436], 1 mM phenylmethylsulfonyl fluoride [Thermo Scientific, 36978]). Roughly 500 µL of acid-washed glass beads (Scientific Instrument Services, GB05) were added, and cells were disrupted for 45 min at 4°C. The cell lysates were collected by centrifugation at 157 x g for 5 min, and the beads were washed once with 500 µL of FA-PIC, resulting in a total volume of 1 mL of cell lysate in FA-PIC. Collected cell lysates were sonicated at 4°C via a Branson Sonifier set at 60% Duty Cycle with an Output Control of 2. Each sample was sonicated on ice for 12 cycles, each consisting of 30 s of sonication and a 30 s period on ice. Chromatin was isolated by centrifugation at 4°C for 30 min at 21000 g, and the resulting pellet was discarded. A 50-µL aliquot was then used to prepare input DNA, confirming the chromatin fragmentation to 200-300 base pairs.

### Chromatin Immunoprecipitation (ChIP)

ChIP was performed as described previously [86]. Briefly, 60 µL of anti-mouse (ThermoFisher, 11040) or 40 µL of anti-rabbit (ThermoFisher, 11204) magnetic Dynabeads were washed twice with chilled PBS (KD Medical, RGF-3200) in BSA (5 mg/mL; Millipore Sigma, A3294)), then incubated for 3 h at 4°C in 200 µL PBS in BSA with 3 µL antibodies (Rpb3: Biolegends, 1YI26l; H3: Abcam, Ab1791, and H4Ac: Sigma, 06-866). After washing the beads, they were incubated with 80 µL of chromatin in 60 µL PBS in BSA and 40 µL of fresh FA-PIC for 3 h (for Rpb3 and H3) or overnight (for H4Ac), rotating at 4°C. Beads were washed sequentially with chilled PBS in BSA, FA- PIC, wash buffer II (50 mM HEPES-KOH, pH 7.5, 500 mM NaCl, 1 mM EDTA, 0.1% sodium deoxycholate, 1% Triton X-100), wash buffer III (10 mM Tris-HCl, pH 8.0, 250 mM LiCl, 1 mM EDTA, 0.5% sodium deoxycholate, 0.5% NP-40 [Millipore Sigma, 74385]), and water before elution. Chromatin was eluted in 100 µL of elution buffer I (50 mM Tris-HCl, pH 8.0, 10 mM EDTA, 1% SDS) by vortexing for 10 s, incubating the sample in a 65°C water bath for 15 min, and then collecting the supernatant by centrifugation at 17000 g for 10 s. The chromatin was eluted again from the beads with elution buffer II (10 mM Tris-HCl, pH 8.0, 1 mM EDTA, 0.67% SDS). Eluents were combined and reverse crosslinked overnight at 65°C. The samples were treated with 5 µL proteinase K (20 mg/mL; Ambion, AM2548) for 2 h at 37°C. Then, DNA from the samples was extracted twice with chloroform, precipitated in ethanol overnight at -80°C, and resuspended in 50 µL RNase A (Millipore Sigma, 11579681) in TE (10 µg/mL) before purification and library prep.

### Library Preparation and Sequencing

ChIP DNA (5-10 ng) was processed using the NEBNext Ultra II Library prep kit from Illumina (E7465S), and IDT xGen UDT-UMI adapters (IDT, 10005903) to generate adapter-barcoded DNA for sequencing as described previously [87]. Libraries were amplified according to the manufacturer’s instructions (9-16 cycles). DNA libraries were bead- and gel-purified prior to paired-end sequencing on a NovaSeq X Plus at Novogene, USA.

### Data Analysis

Sequence reads were trimmed to remove adapters using Cutadapt (parameter: - m 20) and were aligned to the SacCer3 genome using Bowtie2 (parameters: -X 1000- very-sensitive-no-mixed-no-unal). PCR duplicates were removed from .bam files using Samtools. The .bam files were analyzed using BamR (https://github.com/rchereji/bamR). ChIP-seq reads were normalized to a value of 1 for each chromosome. The metagene profiles, scatterplots, and boxplots were generated using JMP. Venn diagrams were generated using BioVenn (https://www.biovenn.nl/). The KEGG pathway analyses were performed at http://bioinformatics.sdstate.edu/go/.

## Abbreviations

AA: amino acid;
ATG: autophagy related;
ChIP-seq: chromatin immunoprecipitation coupled with sequencing;
H4Ac: acetylated histone 4;
HAT: histone acetyltransferase;
HDAC: histone deacetylase;
-N: minus nitrogen/nitrogen starvation;
NDR: nucleosome- depleted region;
Nuc: nucleosome;
ORF: open reading frame;
Pol II: RNA polymerase II;
RiBi: ribosomal biogenesis;
RP: ribosomal protein;
RPG: ribosomal protein gene;
SM: sulfometuron methyl;
TSS: transcription start site;
YPD: yeast extract-peptone- dextrose/nutrient-rich medium

## DATA AVAILABILITY STATEMENT

The accession numbers for the raw and analyzed data reported in this paper are GSE330872. The strains are available upon request.

## ACKNOWLEDGEMENT

We are grateful to the members of the Govind and Klionsky labs for providing comments on the manuscript.

---

**Figure S1.**
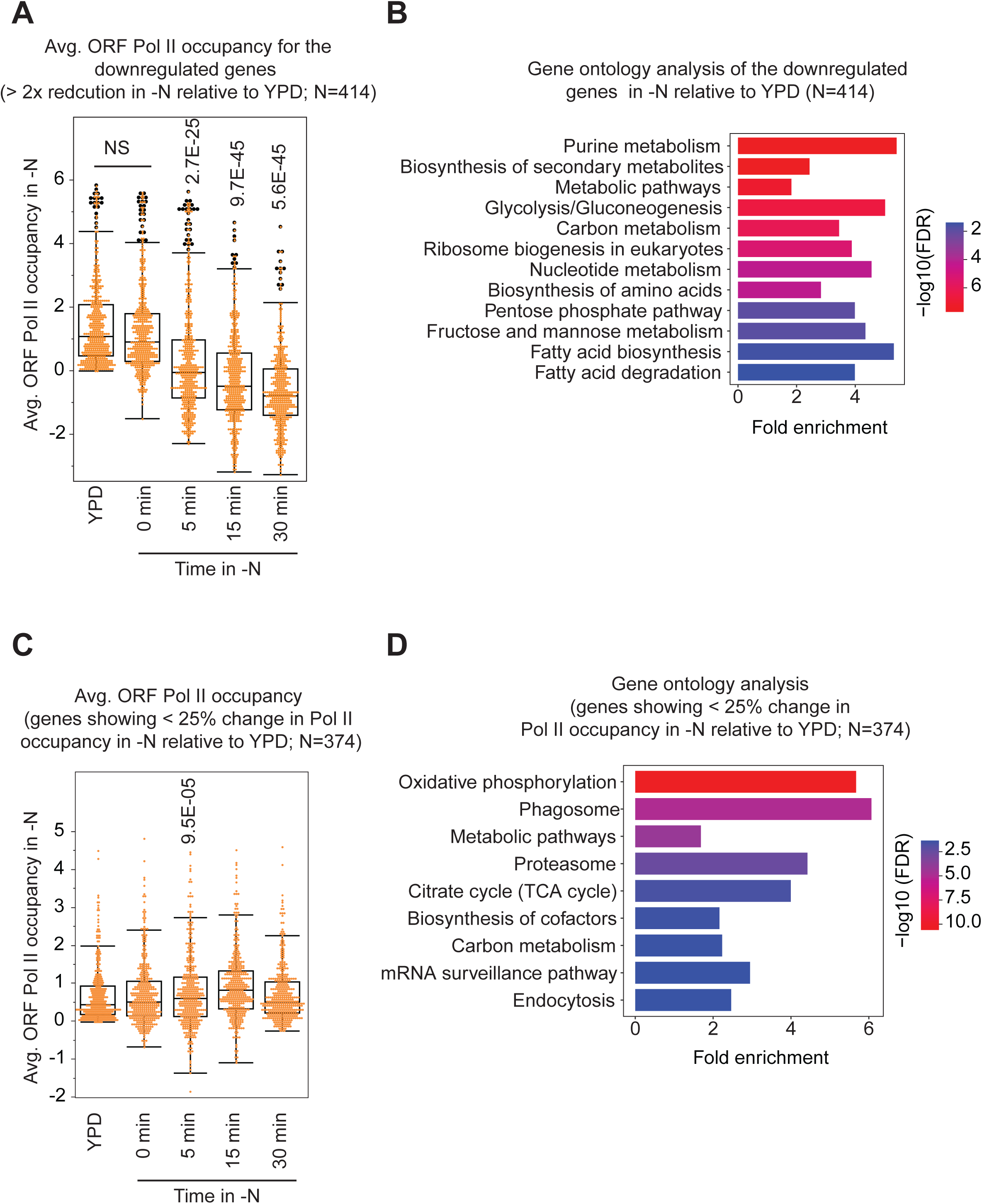
Transcription of a subset of genes is maintained throughout nitrogen starvation. (**A and B**). Box plot showing Pol II occupancies for downregulated genes in YPD and at different time points during nitrogen starvation (A). Each dot represents a gene. These genes were among the top 25% of transcribed genes in nutrient-rich conditions and showed at least a 2-fold reduction in ORF Pol II occupancy in -N medium within 30 min of starvation. The gene-set enrichment for these genes is shown (B). (**C and D**) The box plot shows Pol II occupancies for genes that exhibit less than a 25% change between -N and YPD (C). The gene-set enrichment for these genes is shown (D). False discovery rates (FDRs) are computed via the Benjamini-Hochberg method to correct for multiple testing. Fold enrichment is defined as the percentage of genes in a pathway divided by the corresponding percentage in the background. Related to Figure 1.

**Figure S2.**
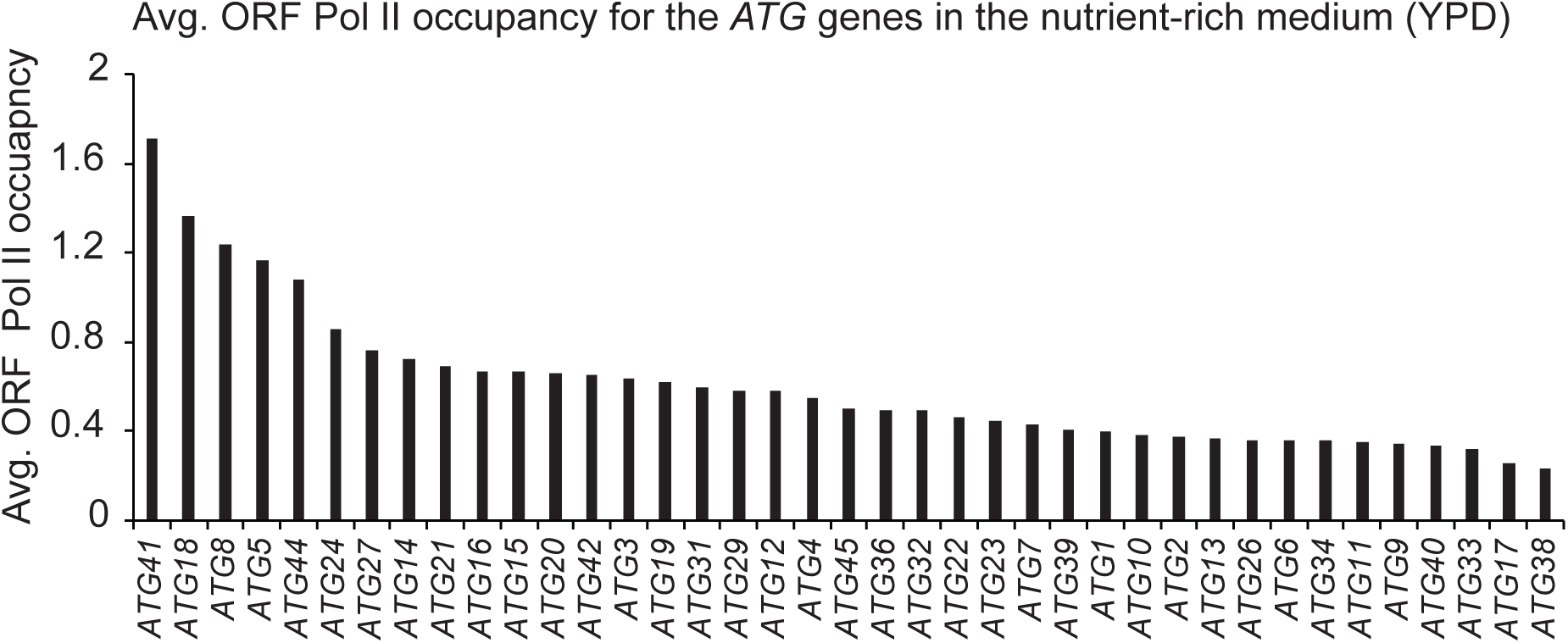
A small subset of genes is expressed at high levels in nutrient-rich medium. Average Pol II occupancies after normalization are shown for *ATG* genes in nutrient-rich medium (YPD). The genes are ordered based on the Pol II occupancy in their ORFs. Related to Figure 2.

**Figure S3.**
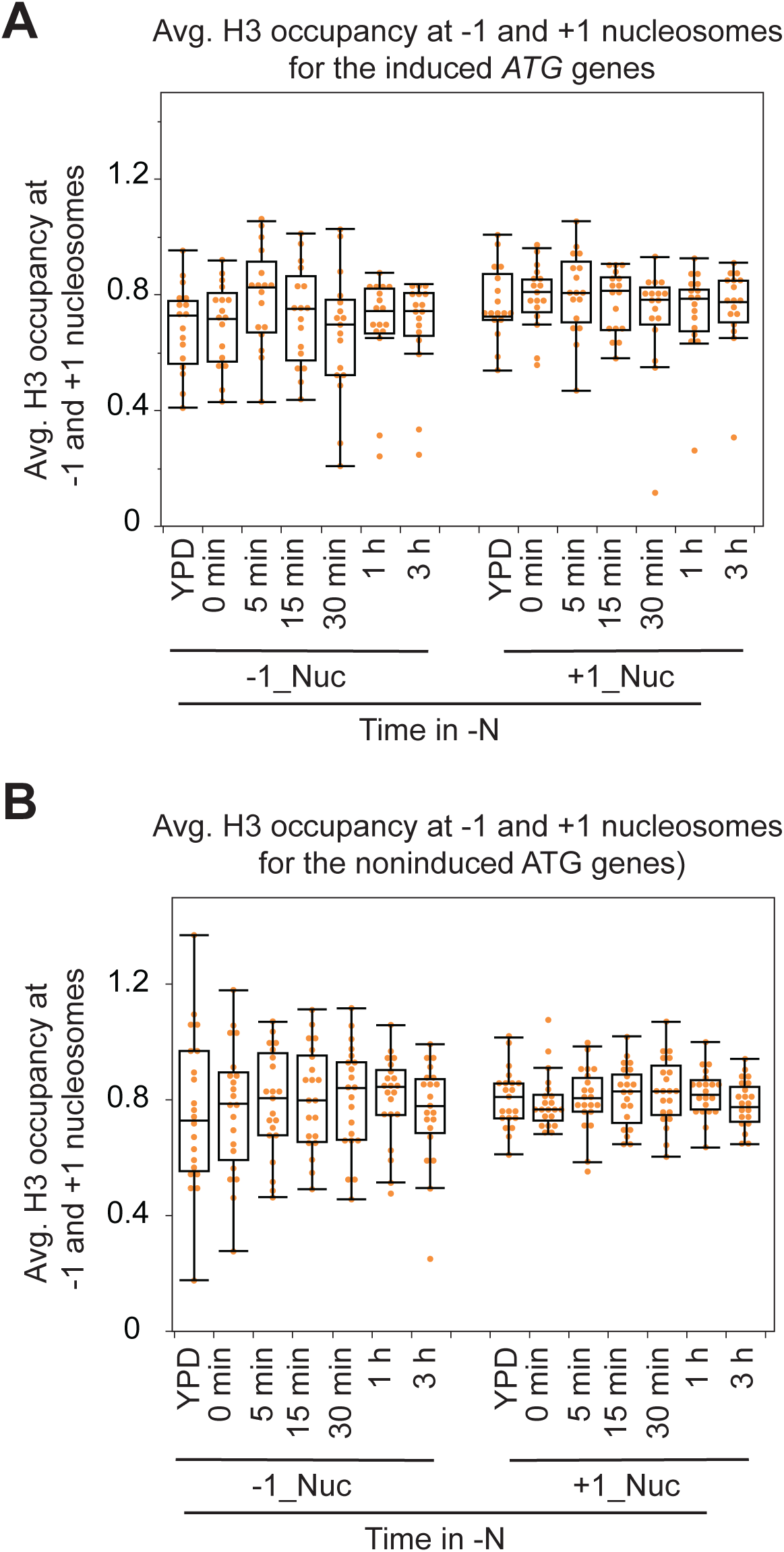
*ATG* genes do not display significant H3 changes in N-starvation conditions. (**A-B**) Normalized H3 ChIP-seq occupancies during N starvation at the induced *ATGs* (A) and non-induced *ATGs* (B). Related to Figure 6.

**Figure S4.**
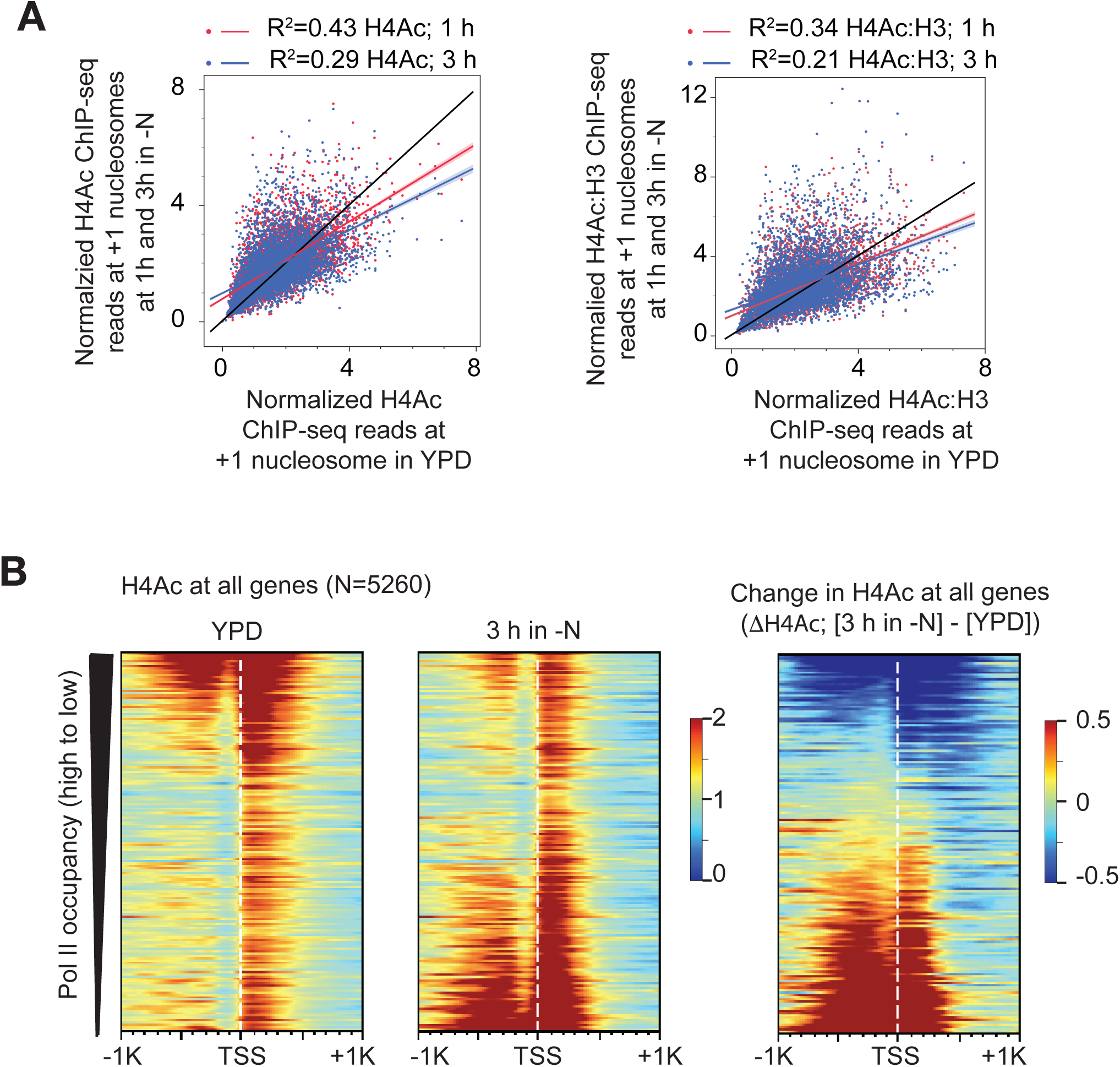
H4 acetylation is reduced in nitrogen starvation. (**A**) H4Ac in nitrogen starvation at 1 and 3 h is plotted against H4Ac (+1 nucleosome position) in YPD for all yeast genes (N=5260) (left panel) and H4Ac:H3 (right panel). (**B**) The H4Ac levels in YPD (left panel), H4Ac under nitrogen starvation for 3 h (middle panel), and H4Ac (-N 3 h minus YPD) are depicted as heatmaps. The genes are ordered from highest to lowest Pol II occupancy.

**Table S1.**
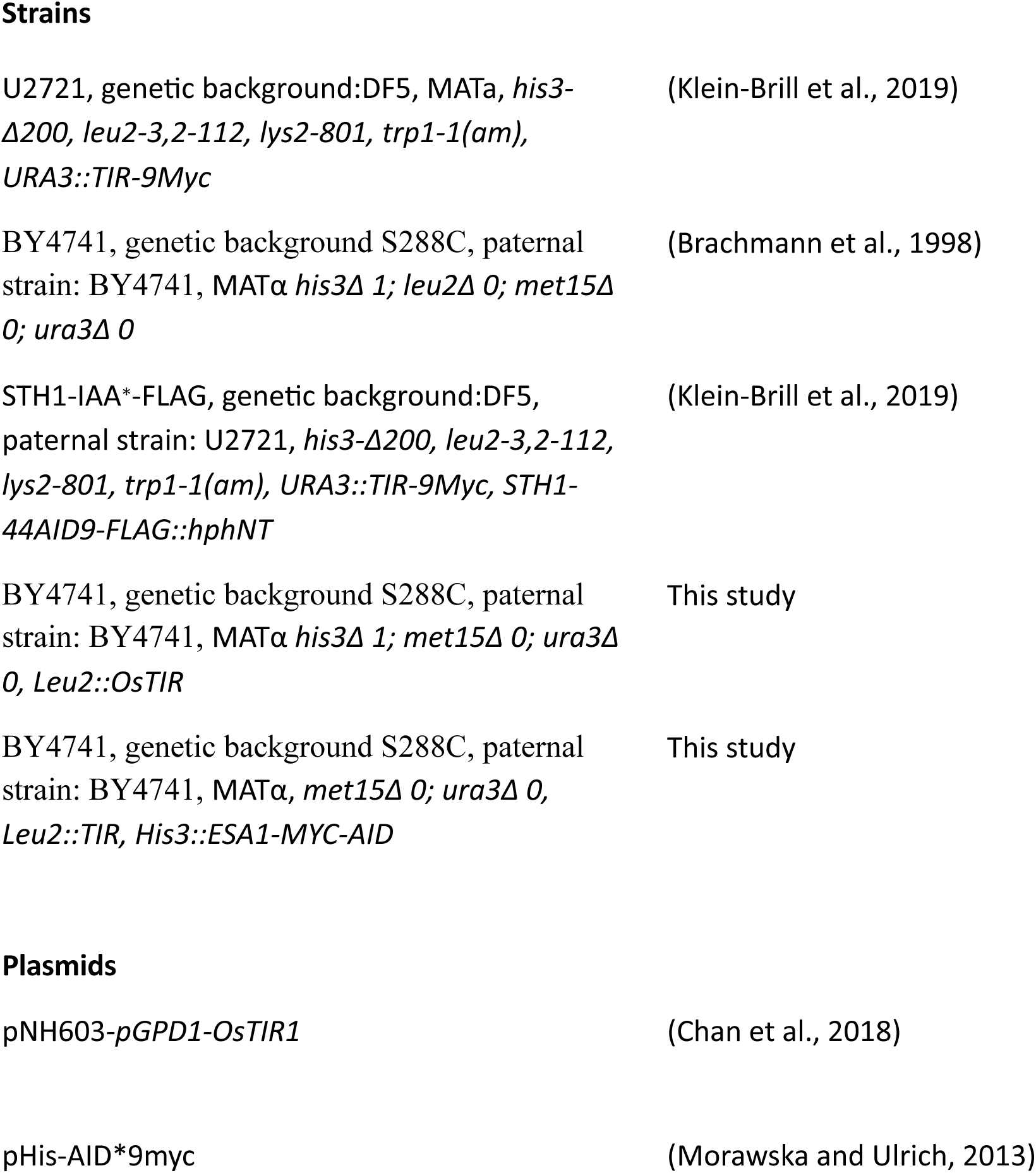
*S. cerevisiae* strains and plasmids.

## References

1. Lei, Y. and D.J. Klionsky, Transcriptional regulation of autophagy and its implications in human disease. Cell Death & Differentiation, 2023. 30(6): p. 1416–1429.

2. Guo, J.Y. and E. White, Autophagy, Metabolism, and Cancer. Cold Spring Harb Symp Quant Biol, 2016. 81: p. 73–78.

3. Ohsumi, Y., Molecular mechanism of autophagy in yeast, Saccharomyces cerevisiae. Philos Trans R Soc Lond B Biol Sci, 1999. 354(1389): p. 1577–1580.

4. Takeshige, K., et al., Autophagy in yeast demonstrated with proteinase-deficient mutants and conditions for its induction. J Cell Biol, 1992. 119(2): p. 301–11.

5. Ohsumi, Y., Historical landmarks of autophagy research. Cell Res, 2014. 24(1): p. 9–23.

6. Klionsky, D.J. and S.D. Emr, Autophagy as a regulated pathway of cellular degradation. Science, 2000. 290: p. 1717–1721.

7. Kraft, C., et al., Mature ribosomes are selectively degraded upon starvation by an autophagy pathway requiring the Ubp3p/Bre5p ubiquitin protease. Nat Cell Biol, 2008. 10(5): p. 602–10.

8. Mao, K. and D.J. Klionsky, Participation of mitochondrial fission during mitophagy. Cell Cycle, 2013. 12(19): p. 3131–2.

9. Huang, Y. and D.J. Klionsky, Identification of the YIPF3-YIPF4 heterodimer as a novel Golgiphagy receptor. Autophagy, 2024: p. 1–2.

10. Papinski, D., et al., Early steps in autophagy depend on direct phosphorylation of Atg9 by the Atg1 kinase. Mol Cell, 2014. 53(3): p. 471–83.

11. White, E., J.M. Mehnert, and C.S. Chan, Autophagy, Metabolism, and Cancer. Clin Cancer Res, 2015. 21(22): p. 5037–46.

12. Cutler, N.S., J. Heitman, and M.E. Cardenas, TOR kinase homologs function in a signal transduction pathway that is conserved from yeast to mammals. Mol Cell Endocrinol, 1999. 155: p. 135–142.

13. Klionsky, D.J., Autophagy: from phenomenology to molecular understanding in less than a decade. Nat Rev Mol Cell Biol, 2007. 8(11): p. 931–7.

14. Feng, Y., Z. Yao, and D.J. Klionsky, How to control self-digestion: transcriptional, post-transcriptional, and post-translational regulation of autophagy. Trends Cell Biol, 2015. 25(6): p. 354–63.

15. Delorme-Axford, E. and D.J. Klionsky, Transcriptional and post-transcriptional regulation of autophagy in the yeast Saccharomyces cerevisiae. J Biol Chem, 2018. 293(15): p. 5396–5403.

16. Delorme-Axford, E., X. Wen, and D.J. Klionsky, The yeast transcription factor Stb5 acts as a negative regulator of autophagy by modulating cellular metabolism. Autophagy, 2023. 19(10): p. 2719–2732.

17. Scott, S.V., et al., Cvt19 is a receptor for the cytoplasm-to-vacuole targeting pathway. Mol Cell, 2001. 7(6): p. 1131–41.

18. Kanki, T., et al., A genomic screen for yeast mutants defective in selective mitochondria autophagy. Mol Biol Cell, 2009. 20(22): p. 4730–8.

19. Mochida, K., Y. Ohsumi, and H. Nakatogawa, Hrr25 phosphorylates the autophagic receptor Atg34 to promote vacuolar transport of α-mannosidase under nitrogen starvation conditions. FEBS Lett, 2014. 588(21): p. 3862–9.

20. Mochida, K., et al., Receptor-mediated selective autophagy degrades the endoplasmic reticulum and the nucleus. Nature, 2015. 522(7556): p. 359–62.

21. Bartholomew, C.R., et al., Ume6 transcription factor is part of a signaling cascade that regulates autophagy. Proceedings of the National Academy of Sciences, 2012. 109(28): p. 11206–11210.

22. Govind, C.K., et al., Phosphorylated Pol II CTD recruits multiple HDACs, including Rpd3C(S), for methylation-dependent deacetylation of ORF nucleosomes. Mol Cell, 2010. 39(2): p. 234–46.

23. Kremer, S.B. and D.S. Gross, SAGA and Rpd3 Chromatin Modification Complexes Dynamically Regulate Heat Shock Gene Structure and Expression. Journal of Biological Chemistry, 2009. 284:: p. 32914–32931.

24. Kurdistani, S.K., et al., Genome-wide binding map of the histone deacetylase Rpd3 in yeast. Nat Genet, 2002. 31(3): p. 248–254.

25. Jin, M., et al., Transcriptional regulation by Pho23 modulates the frequency of autophagosome formation. Curr Biol, 2014. 24(12): p. 1314–1322.

26. Li, X., et al., The TORC1 activates Rpd3L complex to deacetylate Ino80 and H2A.Z and repress autophagy. Sci Adv, 2023. 9(10): p. eade8312.

27. Yi, C., et al., Function and molecular mechanism of acetylation in autophagy regulation. Science, 2012. 336(6080): p. 474–7.

28. Ginsburg, D.S., C.K. Govind, and A.G. Hinnebusch, NuA4 Lysine Acetyltransferase Esa1 Is Targeted to Coding Regions and Stimulates Transcription Elongation with Gcn5. Mol. Cell. Biol., 2009. 29(24): p. 6473–6487.

29. Clarke, A.S., et al., Esa1p Is an Essential Histone Acetyltransferase Required for Cell Cycle Progression. Molecular and Cellular Biology, 1999: p. 2515–2526.

30. Smith, E.R., et al., ESA1 is a histone Acetyltransferase that is essential for growth in yeast. Proc Natl Acad Sci U S A, 1998. 95: p. 3561–3565.

31. Reid, J.L., et al., Coordinate regulation of yeast ribosomal protein genes is associated with targeted recruitment of Esa1 histone acetylase. Mol Cell, 2000. 6(6): p. 1297–1307.

32. Bruzzone, M.J., et al., Distinct patterns of histone acetyltransferase and Mediator deployment at yeast protein-coding genes. Genes & Development, 2018. 32(17-18): p. 1252–1265.

33. Bernard, A., et al., A large-scale analysis of autophagy-related gene expression identifies new regulators of autophagy. Autophagy, 2015. 11(11): p. 2114–2122.

34. Onodera, J. and Y. Ohsumi, Autophagy is required for maintenance of amino acid levels and protein synthesis under nitrogen starvation. J Biol Chem, 2005. 280(36): p. 31582–6.

35. Liu, K., B.M. Sutter, and B.P. Tu, Autophagy sustains glutamate and aspartate synthesis in Saccharomyces cerevisiae during nitrogen starvation. Nature Communications, 2021. 12(1): p. 57.

36. Hardwick, J.S., et al., Rapamycin-modulated transcription defines the subset of nutrient-sensitive signaling pathways directly controlled by the Tor proteins. Proc Natl Acad Sci U S A, 1999. 96(26): p. 14866–70.

37. Pan, X. and J. Heitman, Cyclic AMP-dependent protein kinase regulates pseudohyphal differentiation in Saccharomyces cerevisiae. Mol Cell Biol, 1999. 19(7): p. 4874–87.

38. Young, J.C., et al., Pathways of chaperone-mediated protein folding in the cytosol. Nature Reviews Molecular Cell Biology, 2004. 5(10): p. 781–791.

39. Russo, P., et al., Dual regulation by heat and nutrient stress of the yeast HSP150 gene encoding a secretory glycoprotein. Molecular and General Genetics MGG, 1993. 239(1): p. 273–280.

40. Scott, S.V., et al., Cytoplasm-to-vacuole targeting and autophagy employ the same machinery to deliver proteins to the yeast vacuole. Proc Natl Acad Sci U S A, 1996. 93(22): p. 12304–8.

41. Guan, J., et al., Cvt18/Gsa12 is required for cytoplasm-to-vacuole transport, pexophagy, and autophagy in Saccharomyces cerevisiae and Pichia pastoris. Mol Biol Cell, 2001. 12(12): p. 3821–38.

42. Cheong, H., et al., The Atg1 kinase complex is involved in the regulation of protein recruitment to initiate sequestering vesicle formation for nonspecific autophagy in Saccharomyces cerevisiae. Mol Biol Cell, 2008. 19(2): p. 668–81.

43. Yao, Z., et al., Atg41/Icy2 regulates autophagosome formation. Autophagy, 2015. 11(12): p. 2288–99.

44. Huang, W.P., et al., The itinerary of a vesicle component, Aut7p/Cvt5p, terminates in the yeast vacuole via the autophagy/Cvt pathways. J Biol Chem, 2000. 275(8): p. 5845–51.

45. Shore, D., S. Zencir, and B. Albert, Transcriptional control of ribosome biogenesis in yeast: links to growth and stress signals. Biochem Soc Trans, 2021. 49(4): p. 1589–1599.

46. De Filippi, L., et al., Membrane stress is coupled to a rapid translational control of gene expression in chlorpromazine-treated cells. Curr Genet, 2007. 52(3-4): p. 171–185.

47. Jorgensen, P., et al., A dynamic transcriptional network communicates growth potential to ribosome synthesis and critical cell size. Genes & Development, 2004. 18(20): p. 2491–2505.

48. Rødkær, S.V. and N.J. Færgeman, Glucose- and nitrogen sensing and regulatory mechanisms in Saccharomyces cerevisiae. FEMS Yeast Research, 2014. 14(5): p. 683–696.

49. Powers, T. and P. Walter, Regulation of ribosome biogenesis by the rapamycin-sensitive TOR-signaling pathway in Saccharomyces cerevisiae. Mol Biol Cell, 1999. 10(4): p. 987–1000.

50. Govind, C.K., et al., Simultaneous recruitment of coactivators by Gcn4p stimulates multiple steps of transcription in vivo. Mol Cell Biol, 2005. 25(13): p. 5626–5638.

51. Rawal, Y., et al., SWI/SNF and RSC cooperate to reposition and evict promoter nucleosomes at highly expressed genes in yeast. Genes Dev, 2018. 32(9-10): p. 695–710.

52. Natarajan, K., et al., Transcriptional profiling shows that Gcn4p is a master regulator of gene expression during amino acid starvation in yeast. Mol Cell Biol, 2001. 21(13): p. 4347–4368.

53. Hinnebusch, A.G., Evidence for translational regulation of the activator of general amino acid control in yeast. Proc Natl Acad Sci USA, 1984. 81: p. 6442–6446.

54. Huang, T., et al., The RNA polymerase II subunit Rpb9 activates ATG1 transcription and autophagy. EMBO reports, 2022. 23(11): p. e54993.

55. Qiu, H., et al., An array of coactivators is required for optimal recruitment of TATA binding protein and RNA polymerase II by promoter-bound Gcn4p. Mol Cell Biol, 2004. 24(10): p. 4104–4117.

56. Yoon, S., et al., Recruitment of the ArgR/Mcm1p repressor is stimulated by the activator Gcn4p: a self-checking activation mechanism. Proc Natl Acad Sci U S A, 2004. 101(32): p. 11713–11718.

57. Govind, C.K., et al., Gcn5 promotes acetylation, eviction, and methylation of nucleosomes in transcribed coding regions. Mol Cell, 2007. 25(1): p. 31–42.

58. Qiu, H., et al., Genome-wide cooperation by HAT Gcn5, remodeler SWI/SNF, and chaperone Ydj1 in promoter nucleosome eviction and transcriptional activation. Genome Res, 2016. 26(2): p. 211–25.

59. Zheng, Q., et al., Differential requirements for Gcn5 and NuA4 HAT activities in the starvation-induced versus basal transcriptomes. Nucleic Acids Research, 2023. 51(8): p. 3696–3721.

60. Korber, P., et al., Evidence for Histone Eviction in trans upon Induction of the Yeast PHO5 Promoter. Molecular and Cellular Biology, 2004. 24(24): p. 10965–10974.

61. Qiu, H., et al., Chromatin remodeler Ino80C acts independently of H2A.Z to evict promoter nucleosomes and stimulate transcription of highly expressed genes in yeast. Nucleic Acids Res, 2020. 48(15): p. 8408–8430.

62. Qiu, H.F., et al., Genome-wide cooperation by HAT Gcn5, remodeler SWI/SNF, and chaperone Ydj1 in promoter nucleosome eviction and transcriptional activation. Genome Research, 2016. 26(2): p. 211–225.

63. Spain, M.M., et al., The RSC complex localizes to coding sequences to regulate Pol II and histone occupancy. Mol Cell, 2014. 56(5): p. 653–66.

64. Biernat, E., et al., Histone Acetyltransferases Gcn5 and Esa1 Regulate Occupancy of RSC to Maintain Nucleosome-Depleted Regions and Promote RSC Recruitment to Coding Regions Genome-Wide in Saccharomyces cerevisiae. Mol Cell Biol, 2025: p. 1–23.

65. Biernat, E., M. Verma, and C.K. Govind, Genome-wide regulation of Pol II, FACT, and Spt6 occupancies by RSC in Saccharomyces cerevisiae. Gene, 2024. 893: p. 147959.

66. Kubik, S., et al., Sequence-Directed Action of RSC Remodeler and General Regulatory Factors Modulates +1 Nucleosome Position to Facilitate Transcription. Mol Cell, 2018. 71(1): p. 89–102 e5.

67. Kim, T. and S. Buratowski, Dimethylation of H3K4 by Set1 Recruits the Set3 Histone Deacetylase Complex to 5’ Transcribed Regions. Cell, 2009. 137(2): p. 259–272.

68. Keogh, M.C., et al., Cotranscriptional set2 methylation of histone H3 lysine 36 recruits a repressive Rpd3 complex. Cell, 2005. 123(4): p. 593–605.

69. Carrozza, M.J., et al., Histone H3 methylation by Set2 directs deacetylation of coding regions by Rpd3S to suppress spurious intragenic transcription. Cell, 2005. 123(4): p. 581–592.

70. Guillemette, B., et al., Variant histone H2A.Z is globally localized to the promoters of inactive yeast genes and regulates nucleosome positioning. PLoS Biol, 2005. 3(12): p. e384.

71. Hardy, S., et al., The euchromatic and heterochromatic landscapes are shaped by antagonizing effects of transcription on H2A.Z deposition. PLoS Genet, 2009. 5(10): p. e1000687.

72. Clapier, C.R., et al., Mechanisms of action and regulation of ATP-dependent chromatin-remodelling complexes. Nat Rev Mol Cell Biol, 2017. 18(7): p. 407–422.

73. Biernat, E., et al., The RSC complex remodels nucleosomes in transcribed coding sequences and promotes transcription in Saccharomyces cerevisiae. Genetics, 2021. 217(4).

74. Wen, X., et al., The transcription factor Spt4-Spt5 complex regulates the expression of ATG8 and ATG41. Autophagy, 2020. 16(7): p. 1172–1185.

75. Huang, T., et al., The RNA polymerase II subunit Rpb9 activates ATG1 transcription and autophagy. EMBO Rep, 2022. 23(11): p. e54993.

76. An, H., et al., Systematic quantitative analysis of ribosome inventory during nutrient stress. Nature, 2020. 583(7815): p. 303–309.

77. Joo, Y.J., et al., Gcn4p-mediated transcriptional repression of ribosomal protein genes under amino-acid starvation. Embo j, 2011. 30(5): p. 859–72.

78. Lempiäinen, H. and D. Shore, Growth control and ribosome biogenesis. Curr Opin Cell Biol, 2009. 21(6): p. 855–63.

79. Warner, J.R., The economics of ribosome biosynthesis in yeast. Trends Biochem Sci, 1999. 24(11): p. 437–40.

80. Hinnebusch, A.G. and G.R. Fink, The General Control of Amino Acid Biosynthetic Genes in the Yeast Saccharomyces Cerevisia. Critical Reviews in Biochemistry, 1986. 21(3): p. 277–317.

81. Thomas, G. and M.N. Hall, TOR signalling and control of cell growth. Current Opinion in Cell Biology, 1997. 9(6): p. 782–787.

82. Cherkasova, V.A. and A.G. Hinnebusch, Translational control by TOR and TAP42 through dephosphorylation of eIF2alpha kinase GCN2. Genes Dev, 2003. 17(7): p. 859–72.

83. Longtine, M.S., et al., Additonal modules for versatile and economical PCR-based gene deletion and modification in Saccharomyces cerevisiae. Yeast, 1998. 14: p. 953–961.

84. Ito, H., et al., Transformation of intact yeast cells treated with alkali cations. J Bacteriol, 1983. 153(1): p. 163–8.

85. Shetty, A., N.I. Reim, and F. Winston, Auxin-Inducible Degron System for Depletion of Proteins in Saccharomyces cerevisiae. Curr Protoc Mol Biol, 2019. 128(1): p. e104.

86. Govind, C.K., D. Ginsburg, and A.G. Hinnebusch, Measuring Dynamic Changes in Histone Modifications and Nucleosome Density during Activated Transcription in Budding Yeast. Methods Mol Biol, 2012. 833: p. 15–27.

87. Biernat, E., et al., Histone Acetyltransferases Gcn5 and Esa1 Regulate Occupancy of RSC to Maintain Nucleosome-Depleted Regions and Promote RSC Recruitment to Coding Regions Genome-Wide in Saccharomyces cerevisiae. Mol Cell Biol, 2025. 45(12): p. 623–645.

## References

1. Klein-Brill, A., et al., Dynamics of Chromatin and Transcription during Transient Depletion of the RSC Chromatin Remodeling Complex. Cell Rep, 2019. 26(1): p. 279–292.e5.

2. Brachmann, C.B., et al., Designer deletion strains derived from Saccharomyces cerevisiae S288C: a useful set of strains and plasmids for PCR-mediated gene disruption and other applications. Yeast, 1998. 14(2): p. 115–32.

3. Chan, L.Y., et al., Non-invasive measurement of mRNA decay reveals translation initiation as the major determinant of mRNA stability. Elife, 2018. 7.

4. Morawska, M. and H.D. Ulrich, An expanded tool kit for the auxin-inducible degron system in budding yeast. Yeast, 2013. 30(9): p. 341–51.

